# Ubiquitination and Phosphorylation are Independently Required for Epsin-Mediated Internalization of Cargo in *S. cerevisiae*

**DOI:** 10.1101/2020.02.07.939082

**Authors:** Arpita Sen, Wen-Chieh Hsieh, Claudia B. Hanna, Chuan-Chih Hsu, McKeith Pearson, W. Andy Tao, R. Claudio Aguilar

**Affiliations:** Department of Biological Sciences, Purdue University, West Lafayette, IN 47907, USA; Aromyx Inc., 319 N. Bernardo Avenue, Mountain View, CA 94043, USA; Section on Cellular Communication, National Institute of Child Health and Human Development, National Institutes of Health, Bethesda, MD 20892, USA; Department of Biochemistry, Purdue University, West Lafayette, IN 47907, USA

**Keywords:** Endocytosis, internalization, Ena1, epsin, ubiquitin, phosphorylation

## Abstract

It is well-known that in addition to its classical role in protein turnover, ubiquitination is required for a variety of membrane protein sorting events. However, and despite substantial progress in the field, a long-standing question remains: *given that all ubiquitin (Ub) units are identical, how do different elements of the sorting machinery recognize their specific cargoes?*

Here we provide an answer to this question as we discovered a mechanism based on the coincidence detection of lysine ubiquitination and Ser/Thr phosphorylation for the endocytic adaptor epsin to mediate the internalization of the yeast Na^+^ pump Ena1.

Internalization of Ena1-GFP was abolished in double epsin knock-out in *S. cerevisiae* and was rescued by re-introducing either one of the 2 yeast epsins, Ent1 or Ent2 in an UIM (<u>U</u>b Interacting <u>M</u>otif)-dependent manner. Further, our results indicate that ubiquitination of its C-terminal Lys^1090^ is needed for internalization of Ena1 and requires the arrestin-related-trafficking adaptor, Art3.

We determined that in addition to ubiquitination of K^1090^, the presence of a Ser/Thr-rich patch (S^1076^TST^1079^) within Ena1 was also essential for its internalization. Our results suggest that this ST motif is targeted for phosphorylation by casein kinases. Nevertheless, phosphorylation of this S/T patch was not required for ubiquitination. Instead, ubiquitination of K^1090^ and phosphorylation of the ST motif were independently needed for epsin-mediated internalization of Ena1.

We propose a model in which a dual detection mechanism is used by Ub-binding elements of the sorting machinery to differentiate among multiple Ub-cargoes.

---

Endocytosis is a vital cellular process required for multiple cellular functions including nutrient uptake and control of cell membrane composition. This process involves the selective retention of cargo proteins into nascent endocytic sites by the endocytic machinery through the recognition of sequence-encoded motifs or post-translationally added tags.

In addition to its classical role in proteosome-mediated protein degradation, ubiquitin (Ub) plays an important role as post-translational tag for protein sorting into different subcellular compartments. Indeed, the Ub-tag mediates membrane protein internalization and targeting to the lysosomal/vacuolar and recycling compartments (reviewed in (1, 2)).

Since Ub constitutes the main trafficking signal in *Saccharomyces cerevisiae* (reviewed in (3)), studies in yeast have been instrumental in advancing our understanding of the mechanisms operating in Ub-mediated sorting. For example, it has been clearly established that budding yeast utilizes Rsp5, the only yeast HECT domain-containing, E3 ubiquitin ligase, to promote cargo ubiquitination relevant to endocytosis and other vesicle trafficking events (3, 4). Indeed, Rsp5-mediated ubiquitination is required for the internalization of cargoes such as the pheromone receptor Ste2 (5, 6), the uracil transporter Fur4 (7), the general amino acid permease Gap1 (8) and the zinc transporter Zrt1 (9) among others.

Although Rsp5 is capable of directly interacting with PPxY motif-containing substrates *via* its WW domains (10–13), this consensus sequence is not present in all cargoes that undergo Ub-dependent trafficking (4, 14). In those cases, the PPxY motif-containing, Arrestin-Related Trafficking (ARTs) adaptor family specifically bridges cargo proteins to Rsp5 (15–20)

Following ubiquitination, plasma membrane cargoes are recognized by endocytic adaptors, such as epsin and Eps15 (or Ede1 in *S. cerevisiae*), which contain ubiquitin-binding elements like the <u>U</u>b-interacting <u>m</u>otifs (UIMs) or <u>Ub</u>-<u>A</u>ssociated (UBA) domains. As a consequence of these interactions, cargo is incorporated into nascent endocytic sites and trafficked towards degradative vacuolar/lysosomal compartments or recycling routes.

Specifically, eps15 is known to be responsible for the internalization of the AMPA receptor (21) and Cx43 (22) among other ubiquitinated cargoes; while epsin it is involved in the uptake of <u>v</u>ascular <u>e</u>ndothelial <u>g</u>rowth factor receptor <u>2</u> (VEGFR2) (23), notch ligands (24–27), and the epithelial sodium channel ENaC (28, 29).

The different relevance of these adaptors for the internalization of various ubiquitinated cargoes highlights a crucial knowledge gap in the field: *how identical ubiquitin units attached to different cargoes mediate specific, but differential recognition by Ub-binding proteins?* (which in turn is crucial to promote the needed trafficking outcomes).

Here we report the identification of the Na^+^ pump Ena1 (<u>E</u>xitus <u>Na</u>tru <u>1</u>) as the first described yeast epsin-specific cargo. Further, we established that the internalization of this cargo required ubiquitination of lysine^1090^ and as expected, also needed integrity of epsin’s UIMs.

Importantly, our results indicate that epsin-mediated endocytosis of Ena1 independently required phosphorylation of an upstream Ser/Thr-rich motif *and* ubiquitination of Lys^1090^ in the cytoplasmic domain of the transporter. Epsin co-immunoprecipitated an Ena1 mutant that emulated constitutive phosphorylation and ubiquitination.

In summary, our results indicate that the yeast endocytic adaptor epsin acts upon its cargo based on the presence of a phosphorylated Ser/Thr motif upstream of a ubiquitinated-lysine. We speculate that other Ub-binding elements of the sorting system (such as Ede1) also use coincidence detection of determinants (likely different from the ones recognized by epsins) to identify their targets among multiple Ub-cargoes.

## RESULTS

### The yeast Na^+^ pump Ena1 is internalized in an epsin-dependent manner

*S. cerevisiae* is known to use Ub as its main sorting tag and has been instrumental to study the function of endocytic adaptors (3). Therefore, we relied on this organism to investigate the mechanism by which epsin recognizes its specific targets among multiple ubiquitinated-cargoes. However, it was first necessary to identify a yeast membrane protein that would substantially rely on this adaptor for its internalization.

In contrast to mammals, yeast does not have homologs for established epsin-specific cargoes such as Notch-ligands and VEGFR2-like receptor tyrosine-kinases. Although ENaC-sodium channel homologs are not present either, *S. cerevisiae* has sodium pumps such as the <u>E</u>xitus <u>Na</u>tru (Ena) protein family. Pumps and channels are evolutionarily/mechanistically related (30, 31) and their plasma membrane levels have been shown to be controlled by Ub-mediated endocytosis (32, 33). Therefore, we speculated that members of the Ena family (Ena1-5) were suitable candidates to be internalized in an epsin-dependent manner.

Given its relevance among family members (34–36), we focused our studies on the Ena1 paralog (also known as Pmr2) using an *ENA1::GFP* construct under control of the *MET25* methionine-repressible promoter (37, 38). Stability of WT and mutated versions of Ena1-GFP was verified by Western blotting, while also monitoring their cellular distribution pattern in comparison to free GFP and ER-retained transmembrane proteins (Supplemental Fig. 1).

In agreement with previous observations (37, 38), we observed that in wild type (WT) cells Ena1 partitioned between the plasma membrane and intracellular compartments (Fig. 1A). Further, more than 95% of Ena1-GFP intracellular structures were stained by the lipophilic dye FM4-64 (following a 30min chase—see Supplemental Materials), indicating that they are connected to the endocytic pathway (Supplemental Fig. 2).

**Figure 1:**
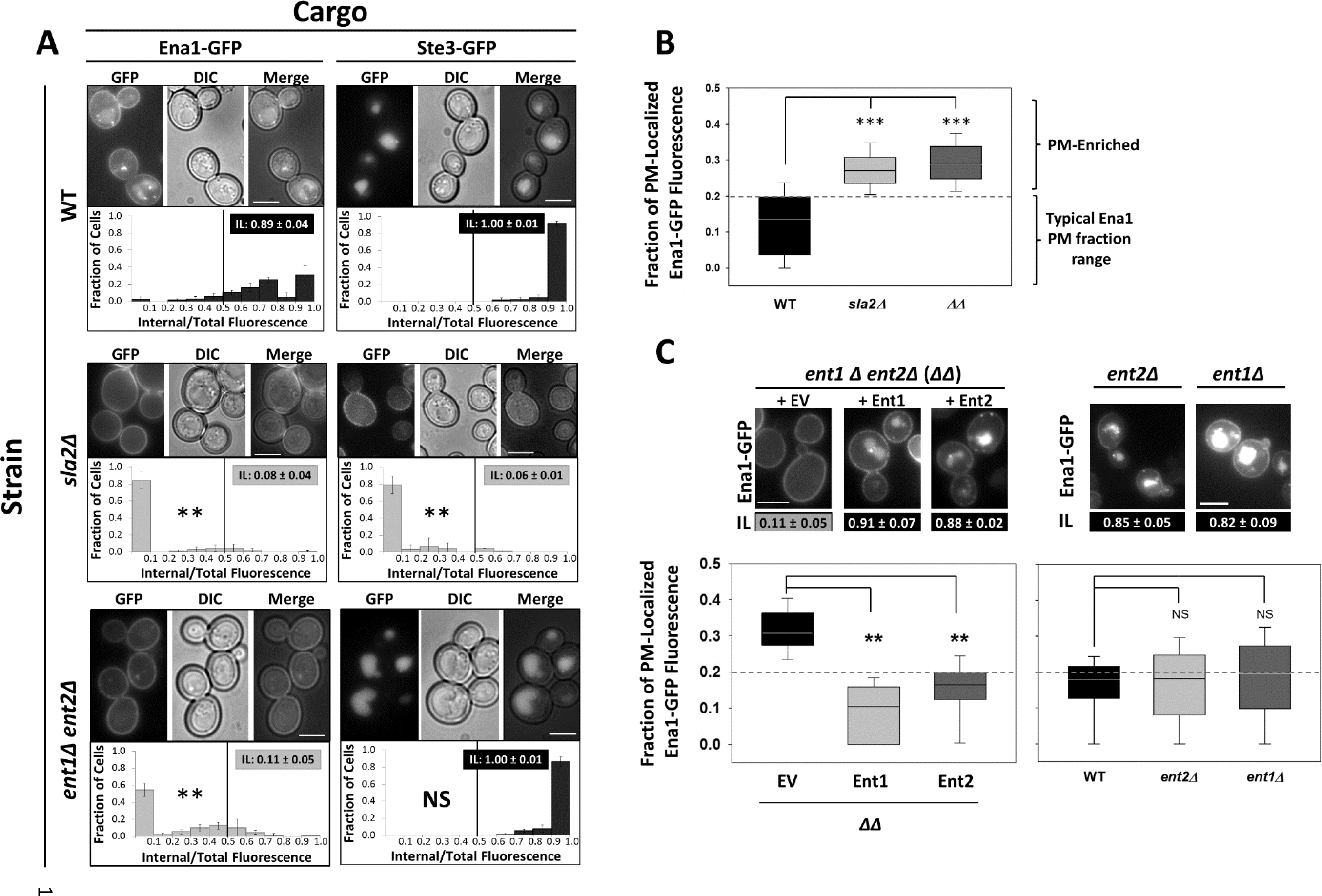
The cellular distribution of Ena1-GFP depends on epsin. **A**. *Localization of Ena1-GFP and Ste3-GFP in wild type (WT), sla2Δ or ent1Δent2Δ (ΔΔ) cells.* Representative images show the cellular distribution of the indicated GFP-tagged cargoes (“GFP”), differential interference contrast (“DIC”) and merged channels. Scale bar, 5μm. Quantification of GFP-fusion protein intracellular localization over 90 cells was performed as described in *Materials and methods*. Results are shown as fraction of cells vs Internal/Total fluorescence histograms; the Kolmogorov-Smirnoff test (with a Bonferroni-corrected α for multiple comparisons αC<0.05/2=0.025) was used to statistically analyze the difference between the distribution of Ena1-GFP (or Ste3-GFP) in WT versus *sla2Δ* and *ΔΔ* cells: **: p< αC; NS: not significant. “Intracellular Localization” (IL) index values (calculated over at least 3 histograms) shown in grey boxes were significantly different from the corresponding WT’s IL (using the Student’s t-test with an α = αC). **B**. *Relative Ena1-GFP plasma membrane (PM) accumulation in WT, sla2Δ or ΔΔ cells.* The fraction of PM-localized Ena1-GFP fluorescence with respect to the total fluorescence (P/T; see *Suppl*. *methods*) in the indicated cell backgrounds are shown as box-plots. The statistical significance of differences between the non-normally distributed P/T values for Ena1-GFP in WT vs *sla2Δ* or *ΔΔ* cells was estimated by using the Wilcoxon’s test with an α = αC. ***: p<< αC. **C**. *Localization and relative Ena1-GFP PM accumulation in cells expressing no epsin or a single paralog.* The values of IL and P/T were estimated for Ena1-GFP expressed in *ΔΔ* cells transformed with empty vector (EV: no epsin present), or with single-copy plasmids carrying either epsin paralog gene from their endogenous promoter; as well as for Ena1-GFP expressed in epsin single KOs (*ent2Δ* and *ent1Δ*: expressing only Ent1 and Ent2, respectively) cells. Representative images are shown.

We quantified the internal GFP-fluorescence intensity as a fraction of the total (I/T) fluorescence intensity and defined an ‘intracellular localization’ (‘IL’) index as the *cumulative fraction of cells* with more than 50% of Ena1 in intracellular compartments (see Supplemental Methods). WT cells exhibited an ‘IL’ of 0.89 ± 0.04 (Fig. 1A) indicating that approximately 89% of this *cell population* contained Ena1 enriched (>50%) in internal compartments. In addition, *individual* WT cells showed Ena1-GFP localized at the plasma membrane (P) corresponding to only13% of their total (T) fluorescence intensity (0.13 P/T median value—Fig. 1B). See *Supplemental methods* for a more detailed description of IL and P/T value estimation and their complementary meaning.

As expected for proteins undergoing endocytosis, Ena1 accumulated at the plasma membrane of the general endocytosis-deficient *sla2Δ* strain and therefore cells have median amount of Ena1-GFP localized at the plasma membrane that doubled as compared to WT cells (Fig. 1B) while exhibiting an IL = 0.08 ± 0.04 (Fig. 1A). As described before (39), internalization of the constitutively endocytosed pheromone receptor Ste3 was also substantially inhibited in *sla2Δ* as compared to WT cells (Fig. 1A, B).

If as hypothesized, Ena1 internalization was dependent on epsin function, then the transporter should be largely localized at the plasma membrane (*i.e.,* should display low IL and high P/T values) in *ent1Δent2Δ* double epsin delete cells (hereafter referred to as ΔΔ). Indeed, as opposed to the WT strain, only ∼11% of the ΔΔ cells had Ena1 enriched in internal compartments (IL = 0.11 ± 0.05—Fig. 1A). Similar to *sla2Δ, ΔΔ* cells showed a substantial accumulation of Ena1-GFP at the plasma membrane (Fig. 1B). In contrast, as expected for a cargo that does not depend on the epsins for internalization (40), Ste3 was properly localized to internal structures in 100% of ΔΔ cells (Fig. 1A). Importantly, re-introduction of either yeast epsin Ent1 or Ent2 could rescue the Ena1 localization defect in ΔΔ cells (Fig. 1C). Along the same lines, Ena1 localization to intracellular compartments was not affected in either single epsin K.O. (*ent1Δ* or *ent2Δ*), also suggesting that the epsins are redundant for promoting Ena1 uptake (Fig.1C).

Further supporting the epsin-specificity of Ena1 internalization, the transporter distribution was not affected by deletion of the genes corresponding to the Yap180 family of endocytic adaptors (data not shown).

Although the Ena1-GFP distribution phenotype in ΔΔ cells was identical to the one shown by the classical endocytosis-defective strain *sla2Δ* (Fig. 1), a direct observation of Ena1 internalization defects in epsin-deficient cells was lacking. Therefore, based on previous knowledge we devised a strategy to directly test the presence of Ena1 endocytosis abnormalities in ΔΔ cells. It is known that under salt stress conditions Ena1 expression is induced and the protein is translocated to the plasma membrane to promote sodium efflux (41–43). Further, Ena1 has been proposed to undergo internalization (44) and vacuolar degradation (45). Therefore, we speculated that upon salt stress *relief,* Ena1 will be subjected to internalization mediated by epsins followed by vacuolar degradation, and that this process will be affected in ΔΔ cells.

To test the potential contribution of epsins to low salt-induced downregulation of Ena1, we designed a salt stress relief assay (see Supplemental Methods). Briefly, WT and ΔΔ cells were subjected to salt stress (0.5M NaCl) following a short induction of Ena1-GFP expression. This was done to ensure that the expressed Ena1-GFP was recruited to and maintained at the plasma membrane to enable the efflux of excess sodium. Under Ena1 expression-suppressed conditions and after lowering the salt concentration in the media, cells were imaged over time and the peripherally-localized fluorescence intensity of Ena1-GFP was quantified at each time-point. Figure 2A shows a representative experiment where at least 50 cells at each time-point were analyzed. In WT cells, the peripheral fluorescence intensity of Ena1-GFP underwent a sharp decrease 45min after salt excess removal, indicating transporter uptake (Fig. 2A-upper panel). In later time points, the vacuolar fluorescence also dramatically diminished, indicating Ena1-GFP degradation (Fig. 2A-upper panel). In striking contrast, the fluorescence intensity of GFP-Ena1 at the periphery of ΔΔ cells remained at high values over time (Fig. 2A-lower panel), indicating that epsins are required for Ena1-GFP removal from the plasma membrane.

**Figure 2:**
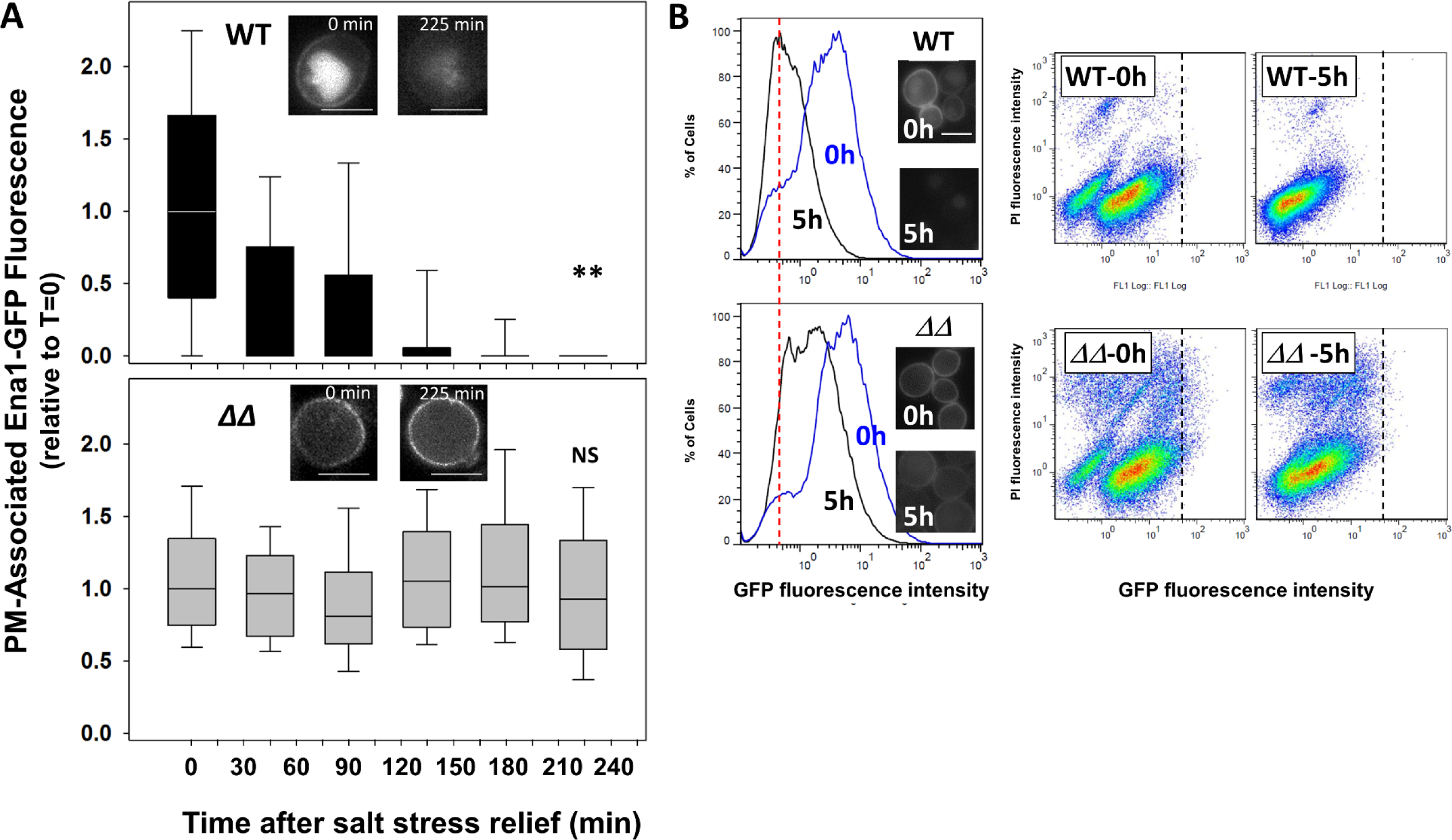
Ena1 internalization is epsin-dependent. **A**. *Salt stress relief stimulates Ena1 internalization in WT, but not ΔΔ cells.* PM-associated fluorescent intensity of Ena1-GFP following salt stress-relief was tracked over time. A representative experiment show PM fluorescence intensity relative to time 0 of >30 cells per time-point. Statistical significance of the difference between the PM-associated fluorescent intensity of Ena1-GFP at time 0 and 225min for WT and *ΔΔ* cells was assesed using Wilcoxon’s test. **: p<0.01; NS: not significant. Representative images for the indicated time-points and for each strain are shown. Scale bar: 5μm. See *Supplemental Methods* for details. **B**. *Removal of Ena1-GFP from the PM is delayed in ΔΔ cells with respect to WT*. Total fluorescent intensity of Ena1-GFP from 5x10^4^ cells was measured by flow cytometry at 0 and 5h after production of Ena1-GFP was halted. % of cells vs GFP-fluorescence intensity histograms are shown for WT (upper left) and *ΔΔ* (lower left) cells at 0 (blue trace) and 5h (black trace). Red vertical dashed line marks the peak position of Ena1-GFP fluorescence intensity in WT cells after 5h-chase; note that the corresponding 5h-peak for *ΔΔ* cells is shifted to the right (higher fluorescence intensity). Representative images of the analyzed cells are shown as insets (scale bar: 10μm). The corresponding Propidium Iodide (PI) vs GFP fluorescence intensity scatter plots are included. Black vertical dashed lines define the GFP fluorescence intensity boundaries at time 0 for WT and *ΔΔ* cells. Note the marked decrease in fluorescence intensity for WT as compared to *ΔΔ* cells after 5h chase. See *Supplemental methods* for details.

The role of epsins in Ena1-GFP retrieval from the plasma membrane was confirmed in complementary experiments using flow cytometry. Specifically, WT and ΔΔ cells were allowed to express Ena1-GFP overnight, and samples were analyzed by flow cytometry at 0h and 5h *under Ena1-GFP expression-suppressed conditions*. Figure 2B shows that 5h after turning off the *MET25* promoter (with 2mM Methionine) fluorescence intensity associated with WT cells was low due to internalization and vacuolar degradation of GFP-Ena1, whereas at 5h, ΔΔ cells displayed high fluorescence intensity closer to the 0h time-point. It should be noted that despite other endocytic adaptors (*e.g.,* Ede1) are present, Ena1 was severely impaired for uptake in ΔΔ cells.

These results indicate that epsins are required for internalization of Ena1 triggered by salt stress relief. Also, as a whole, these experiments support the conclusion that changes in Ena1 cellular distribution (*e.g.,* Fig. 1, and all throughout this study) upon deletion of the PM-localized epsin endocytic adaptors, largely reflects impairment in the Na^+^-transporter uptake.

### Ena1 internalization is mediated by ubiquitination of its C-terminal cytoplasmic segment and requires epsin’s UIMs

Since ubiquitination has been implicated in the trafficking of Ena1 (45) and epsin is a known Ub-binder protein, we hypothesized that the UIMs of epsin were required for Ena1 internalization. Indeed, we observed that Ent1/2 variants bearing mutations that abolish the functionality of the two epsin UIMs (Ent1/2^U^*^m^*: Ent1^S177D, S201D^ and Ent2^S187D, S218D^—(46)) were unable to rescue the Ena1 internalization defect seen in ΔΔ cells (Fig. 3).

**Figure 3:**
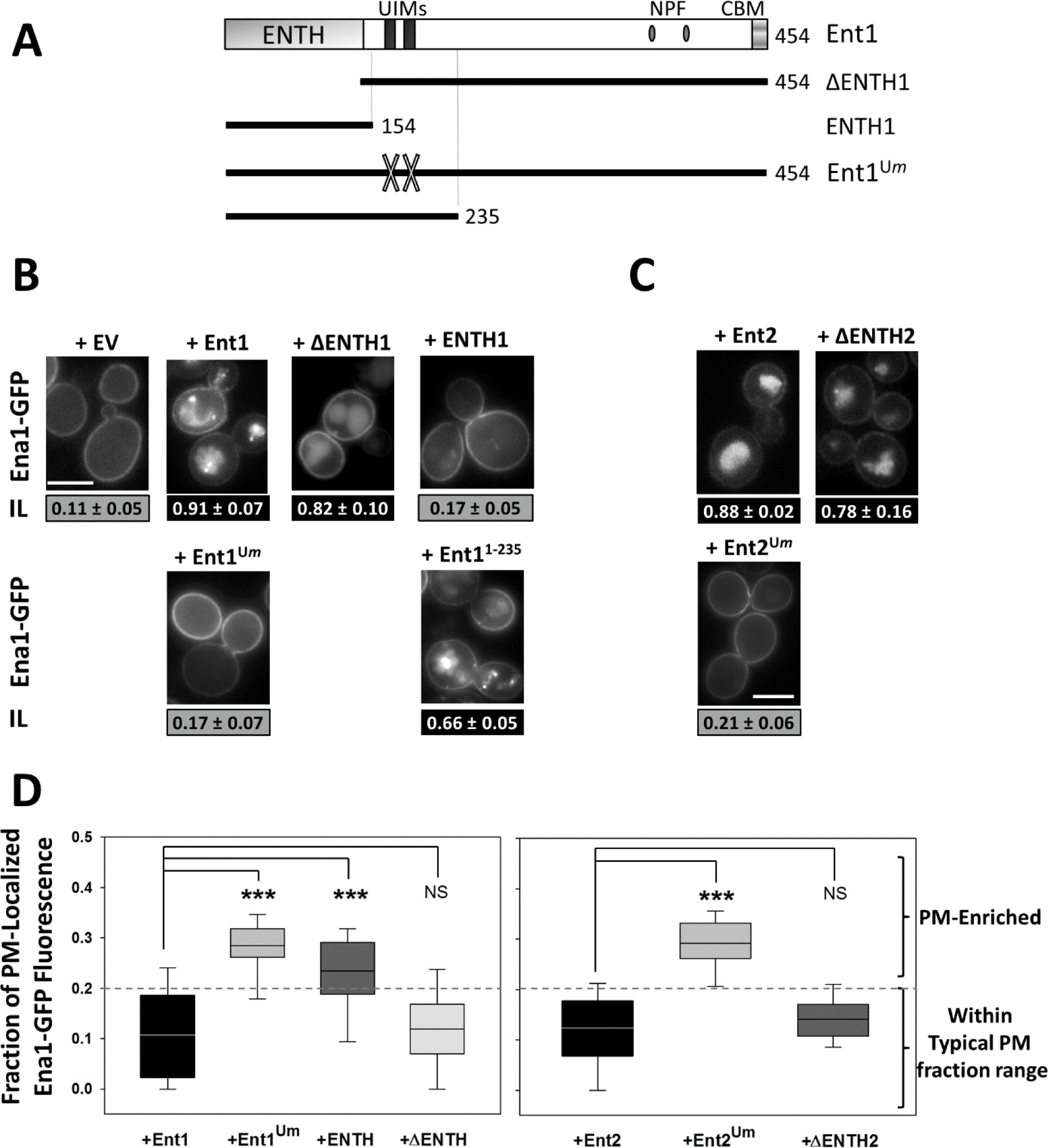
Epsin’s UIMs are required to sustain Ena1 internalization. ΔΔ cells were transformed with empty vector (EV) or plasmids encoding for different Ent1 or Ent2 variants listed in panel **A**. Representative images and internalization indexes for Ent1 (**B**) and Ent2 (**C**) WT and mutated variants are shown. Scale bar: 5μm. ENTH: <u>E</u>psin <u>N</u>-<u>T</u>erminal <u>H</u>omology (domain); NPF: Asn (N)-Pro (P)-Phe (F) tripeptide; CBM: <u>C</u>lathrin <u>B</u>inding <u>M</u>otif; Ent1^U^*^m^*: Ent1 UIM mutant (Ent1^S177D, S201D^); Ent2^U^*^m^*: Ent2 UIM mutant. Image analysis and quantification was performed as described in Fig. 1A. IL values shown in black boxes are statistically different from ΔΔ+EV; (Bonferroni corrected αC<0.05/5=0.01). **D**. Ena1-GFP relative PM accumulation values for cells expressing the indicated Ent1 or Ent2 mutants as sole source of epsin were represented and statistically analyzed (αC=0.01) with respect to Ent1 or Ent2 as in Fig. 1B.

In agreement with these findings, the UIM-containing, C-termini of Ent1 or Ent2 (ΔENTH fragments), but not their lipid-binding ENTH domains, were also sufficient to suppress Ena1 uptake deficiency in ΔΔ cells (Fig. 3). It is worth to reiterate that although ΔΔ cells express other Ub-binding endocytic adaptors such as Ede1, Ena1 uptake showed a strong dependence on epsin UIM integrity.

#### Lys^1090^ is necessary for Ena1 internalization

Since epsin UIMs are required for Ena1 internalization, we hypothesized that this cargo will undergo lysine-ubiquitination as a requisite for endocytosis. It should be noted that Ena1 is a relatively large (1091 residues) protein with 10 predicted transmembrane regions and a large number of cytosol-facing Lys residues as potential targets for ubiquitination. In order to tackle the daunting task of identifying putative internalization-relevant ubiquitination sites, we took into consideration other authors’ pioneering findings working with yeast multi-spanning endocytic cargoes. Specifically, it has been determined that the internalization of the a-factor receptor Ste3, and the α-factor receptor Ste2, required ubiquitination of their cytosolic C-termini (6, 47–49). Based on these findings, we hypothesized that the Ena1 C-terminal tail (residues 1044-1091) is important for its uptake. In agreement with this, C-terminal truncations of Ena1 displayed intracellular distribution defects (Fig. 4A, B). Further, an Ena1 truncation that lacked only the last four amino acids GIKQ (Ena1^Δ1088-1091^) failed to internalize in WT cells (Fig. 4A, B). Since lysines are sites for ubiquitination, we speculated that K^1090^ was the residue within the GIKQ sequence responsible for Ena1 internalization. In fact, the Ena1^K1090R^ mutant was severely impaired for uptake, thereby indicating a requirement of K1090 for Ena1 internalization (Fig. 4C). Point mutations of the adjacent residues (Ena1^G1088A^, Ena1^I1089A^and Ena1^Q1091A^) did not grossly affect the transporter’s localization (although the Q1091A mutant showed a slight accumulation of Ena1 at the PM) (Fig. 4C).

**Figure 4:**
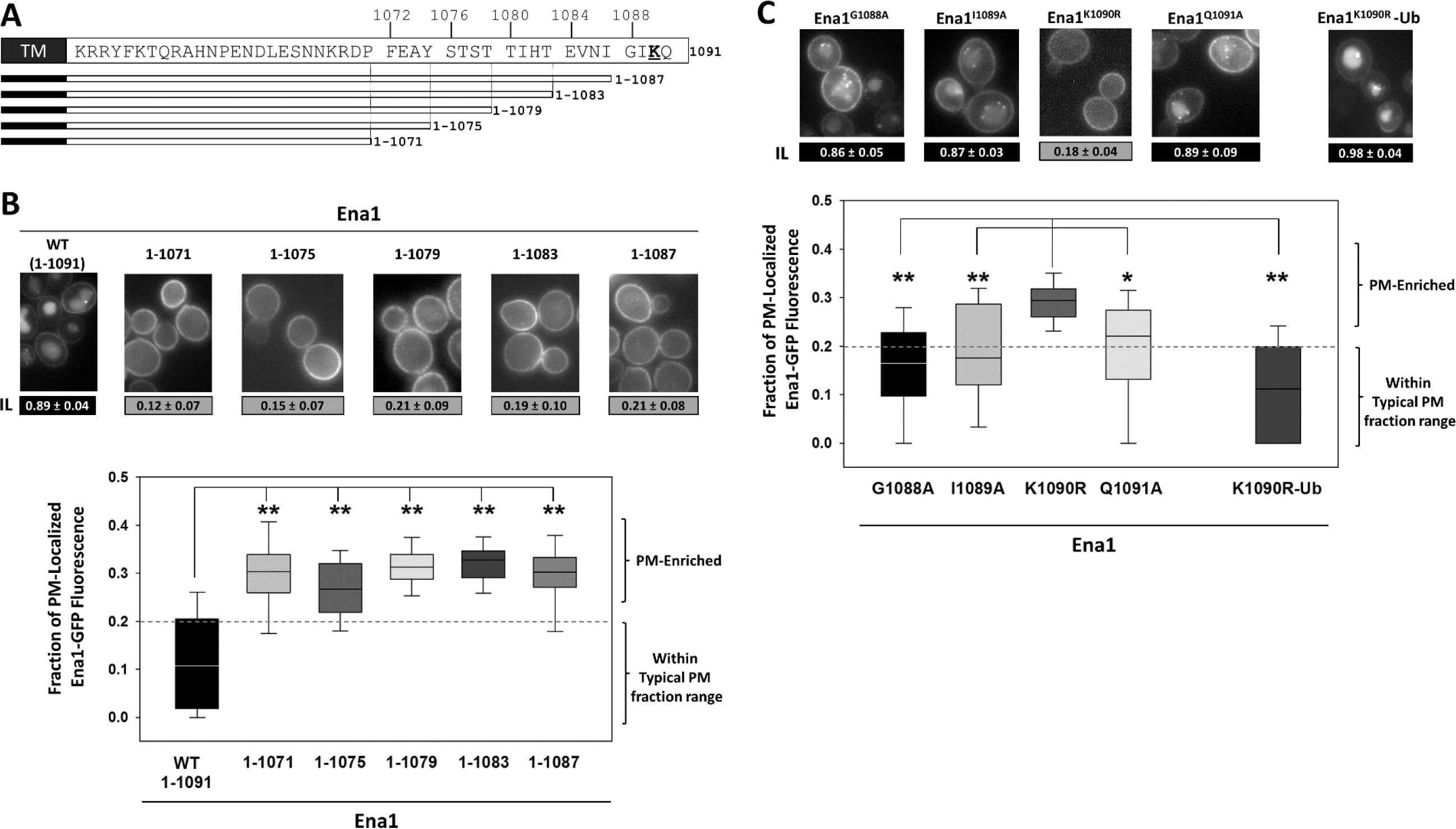
Ena1 K^1090^ is necessary for the transporter internalization. The indicated Ena1-GFP truncations (**A, B**) and point-mutants (**C**) were expressed in yeast cells, IL indexes and relative PM accumulation values (**B**, **C**) were estimated as in Fig. 1. Representative images are included. Scale bar: 5μm. Differences between Ena1^1-1091^ (WT full-length) and truncations (**B**) or between Ena1^K1090R^ and other point mutants or Ena1^K1090R^–Ub (**C**) were statistically analyzed as described in *Suppl. methods* and in Fig. 1 (**: p<<αC=0.01).

Importantly, a C-terminal *in-frame* fusion of ubiquitin with the Ena1^K1090R^ mutant was able to re-establish its intracellular localization (Fig. 4C), strongly suggesting that K^1090^ was a target for ubiquitination.

In addition, we verified that Ena1-Ub fusions used throughout this study were not directly trafficked from Golgi structures to endosomes but were all able to reach (and accumulate at) the plasma membrane in *sla2Δ* cells (data not shown). Experiments performed with *rcy1Δ* (recycling-deficient) strain showed that none of these mutants accumulated in this strain’s characteristic/aberrant compartment suggesting the absence of mistrafficking to recycling routes (data not shown).

#### Ena1 internalization depends on Art3

Rsp5 is the sole E3 ubiquitin ligase in yeast involved in internalization of cargo, which can directly interact with its substrates via its WW domains (3). However, many cargoes do not contain the PPxY motif required for interacting with the WW domains of Rsp5 (4).

Nevertheless, it is very well established that the Arrestin-Related Trafficking (ART) family of adaptors specifically recognizes cargoes and they recruit Rsp5 to mediate selective cargo ubiquitination *via* their own PPxY motif (3, 50). Since Ena1 does not contain a PPxY motif, we tested if its uptake was ART-dependent. We quantified Ena1 localization within a panel of single ART-deleted cells (from *art1Δ* through *art10Δ*—Supplemental Fig. 3). While Ena1 uptake was normal in most of these strains, it was significantly affected in *art3Δ* (Fig. 5A, C) and partially in *art6Δ* cells (Supplemental Fig. 3). It should be noted that Art3 and Art6 are paralogs. Importantly, and as expected, this defect could be bypassed by *in frame*-fusion of ubiquitin (Fig. 5A, C). We also monitored the uptake of Tre1 (Transferrin Receptor-like protein 1), which is a PPxY motif-containing protein that undergoes Rsp5-dependent ubiquitination (51, 52) in an *Art-independent manner*. As predicted, Tre1 internalization was not affected in *art3Δ* cells (Fig. 5A). Importantly, expressing Art3 from a plasmid in *art3Δ* cells re-established Ena1-GFP internalization (Fig. 5B, C).

**Figure 5:**
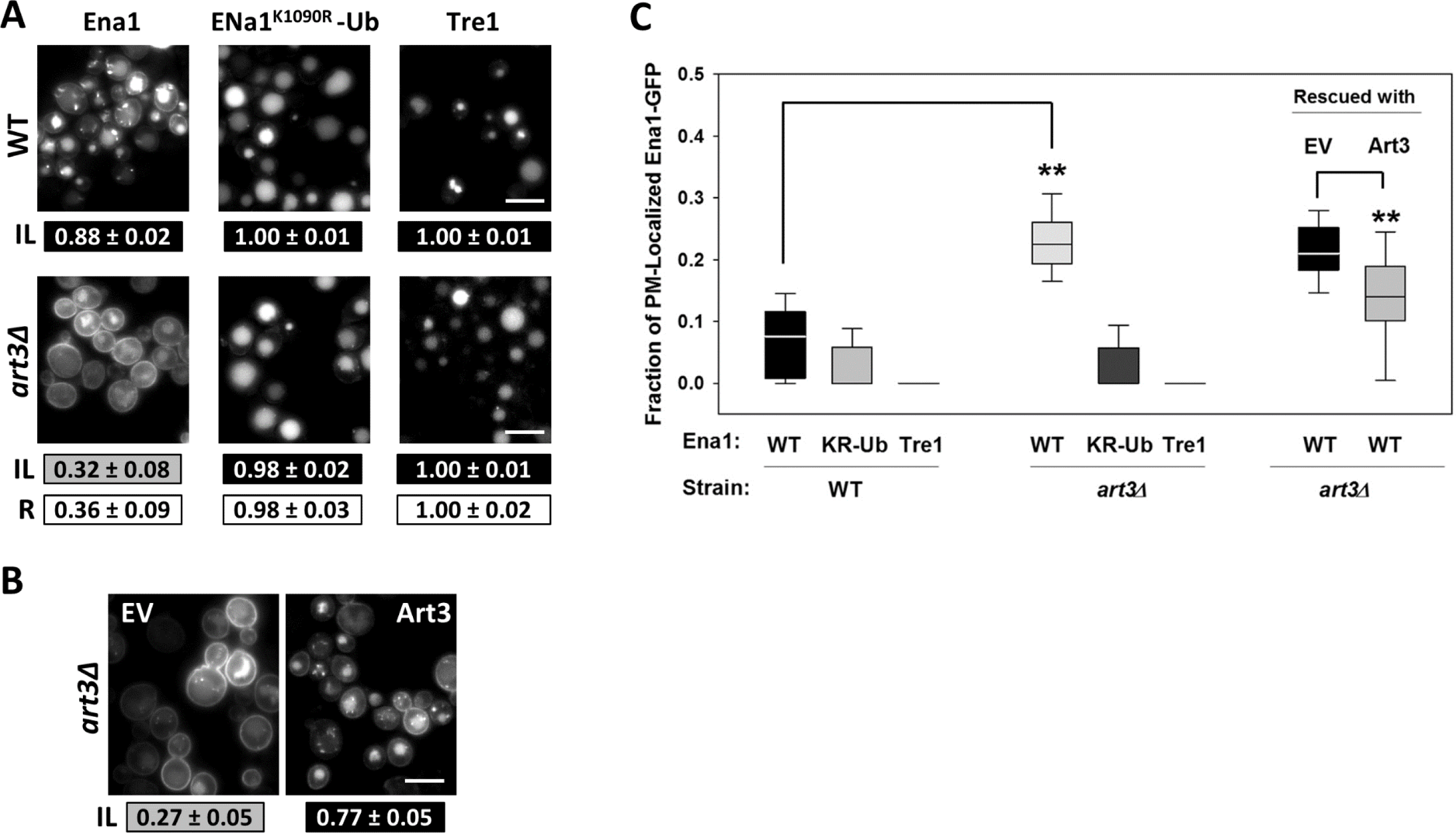
Ena1 internalization is Art3 dependent. **A**. Intracellular localization of Ena1-GFP, Ena1^K1090R^-Ub-GFP and Tre1-GFP was analyzed in WT and *art3Δ* cells. IL indexes were estimated and analyzed as described in Fig. 1. Representative images are included. Scale bar: 10μm. Values in white boxes represent the ratio between the corresponding ILs for each protein in *art3Δ* and WT cells. **B**. Ena1-GFP expressing, *art3Δ* cells were transformed with empty vector (EV) or plasmid for expression of Art3. Cells were imaged and analyzed as described in *Materials and methods*. **C**. Relative PM accumulation for the indicated Ena1 mutants was estimated and statistically analyzed with respect to the corresponding WT strain or EV-transformed *art3Δ* as in Fig. 1. **: p<< α=0.05—Wilcoxon’s test. KR-Ub: Ena1^K1090R^-Ub.

Interestingly, Ena1 uptake was also partially affected by deletion of Art3’s paralog Art6, but not by K.O. of Art9 (Supplemental Fig. 3). Since Art9 has been shown to be involved in Ena1 functional regulation (53), but our data showed that is dispensable for internalization, we speculate that this Art protein might play a role in a different Ena1 trafficking event or other regulatory step.

In summary our results indicate that *Ena1 internalization was impaired*:

-by epsin deletion or epsin UIM mutation (rescued by re-expression of either Ent1 or Ent2 or their corresponding UIM-containing C-terminal fragments).

-by mutation of the C-terminal Ena1 K1090R (rescued by a C-terminal Ub fusion-in-frame).

-by deletion of the Ub-ligase Rsp5 adaptor Art3 (rescued by a C-terminal Ub fusion-in-frame).

Therefore, we speculate that yeast epsins (*via* their UIMs) recognize Ena1 ubiquitinated (by an Art3/Rsp5 complex) at K1090.

### In addition to K^1090^-Ubiquitination, a Ser/Thr-rich region within Ena1 C-terminus is necessary for its internalization

Although our results indicate that Ena1 ubiquitination is necessary for UIM-mediated recognition by epsin, other yeast ubiquitinated cargoes (*e.g.,* Ste3) are internalized independently of Ent1/2. A corollary observation (further emphasizing Ub-binding adaptor specificity) is that even in the presence of other ubiquitin-binding endocytic proteins (*e.g.,* Ede1), Ena1 uptake is severely affected by absence of epsins. Therefore, this led us to a very important unanswered question in the field: given that all Ub units are identical, what are the basis for cargo (*e.g.,* Ena1) specificity for different adaptors (*e.g.,* epsin)?

We speculated that the coincidence detection of ubiquitinated K^1090^, along with another Ena1 putative determinant lead to epsin specificity. Since the C-terminus of Ena1 holds one of such elements (K^1090^), we started our search for a hypothetical Ub-independent determinant within this region. Specifically, we performed a scanning mutational analysis of the Ena1 C-terminal tail by simultaneous mutation of four consecutive amino acids to glycine (Fig. 6). This approach was preferred to alanine scanning to avoid the introduction of a potentially hydrophobic Ala x4 stretch. It should also be highlighted that point mutations or truncations introduced between residues 1050-1070 in Ena1 C-terminus led to ER-retention (see example in Supplemental Fig. 1).

**Figure 6:**
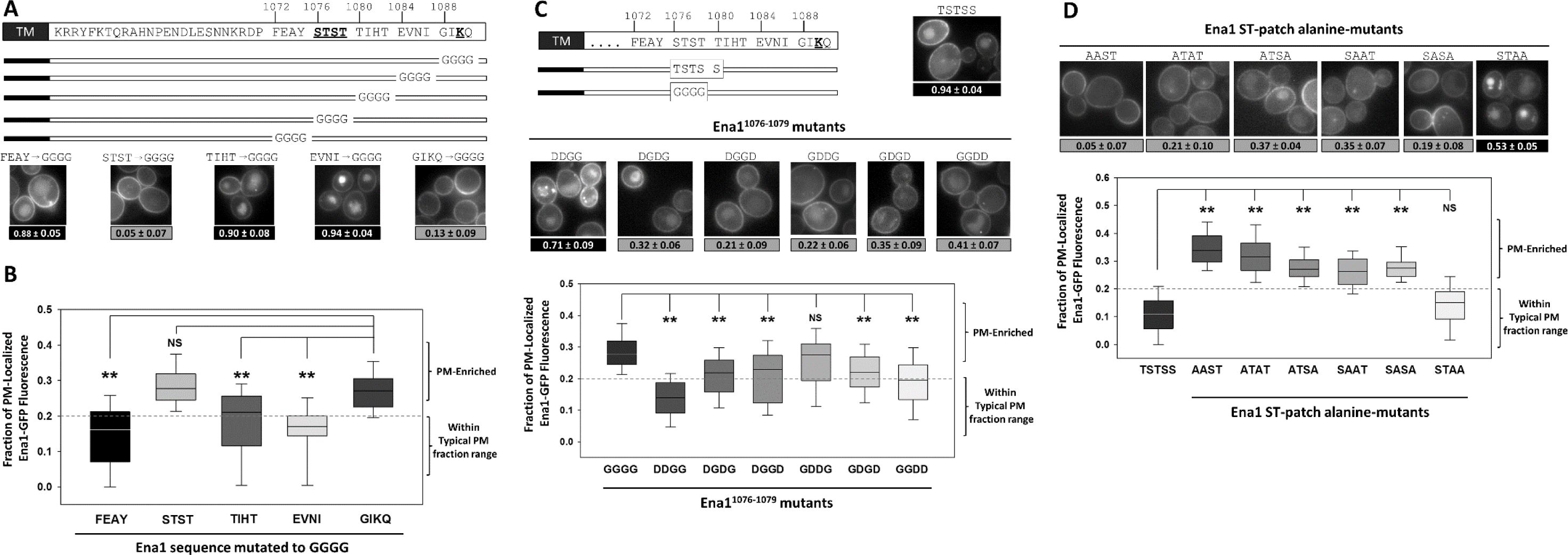
Ena1 S^1076^ T^1077^ residues are required for its internalization and targeted for phosphorylation. The indicated Ena1-GFP mutants were expressed in yeast cells and IL indexes and relative PM accumulation values were estimated as in Fig. 1. Representative images are included. Scale bar: 5μm. Differences between samples were statistically analyzed as described in *Materials and methods* and in Fig. 1 (**: p<<αC=0.008).

As expected, mutation of the G^1088^IKQ^1091^ sequence led to loss of Ena1 internalization by elimination of the critical K^1090^ (Fig. 6A, B). While most Gly x4 Ena1mutants displayed normal endocytosis, mutation of the S^1076^TST^1079^ residues caused a substantial impairment of cargo uptake (Fig. 6A, B). Interestingly, although 90% of cells expressing the T^1080^IHT^1083^-to-GGGG Ena1 mutant showed enrichment of the transporter in intracellular structures (Fig. 6A), they exhibited a PM-associated GFP-fluorescence fraction slightly above normal levels (Fig. 6B). Therefore, we speculate that the Thr residues included in this region are functionally linked to the S^1076^TST^1079^ patch. The identified Ser/Thr-rich (including T^1080^) sequence constituted an obvious candidate for phosphorylation. Indeed, mutational analysis and detection of the corresponding phosphorylated-peptides by mass spectrometry indicated this to be the case (see below).

#### Ena1 C-terminal Ser/Thr patch mutational analysis

Mutation of S^1076^TST^1079^ to TSTS did not affect Ena1 uptake (Fig. 6C, D), suggesting that, rather than the exact sequence, the biochemical nature of Ser/Thr residues (*e.g.,* as targets for phosphorylation) is important for its role in Ena1 internalization. Next, we changed the Gly residues in the uptake-defective G^1076^GGG^1079^ (Ena1^G1076GGG1079^ = Ena1^GGGG^) quadruple mutant to D or E (one at a time or in pairs) to mimic constitutive phosphorylation and to assess the relative importance of the residues within this ST-patch. While single mutants failed, a double D-mutation (Ena1^G1076D, G1077D^ = Ena1^DDGG^) rescued the uptake of Ena1 (Fig. 6C). Similar results were obtained with the equivalent double E mutant Ena1^EEGG^ and Ena1^G1076S, G1077T^ (Ena1^STGG^) (data not shown). All other double D-mutations showed some level of improvement (Fig. 6C): *i.e.,* slight increase of the number of cells with the corresponding Ena1-GFP mutant enriched in intracellular compartments and a decrease in levels of the transporter at the PM as compared to Ena1^GGGG^ (except for the GDDG mutant-Fig. 6C). However, only the Ena1-GFP DDGG and GGDD mutants also showed median P/T values within the normal range for the transporter (Fig. 6C). Since only Ena1^DDGG^ exhibited all signs of substantial rescue (*i.e.,* significant increase in IL values and significant decrease in PM-associated mutant Ena1-GFP levels *compatible with the normal range for the transporter*), we concluded that the first two positions within the ST patch are the most important to mediate Ena1 uptake.

To complement these studies, we generated S/T-to-alanine mutations of the ST-motif in Ena1^WT^ to mimic constitutive dephosphorylation. While all Ena1 ST-mutants displayed defective uptake at certain extent, Ena1^S1076A T1077A^ (Ena1^AAST^) was the most affected (Fig. 6D). In addition, the sole presence of S^1076^ and T^1077^ was enough to maintain proper Ena1-GFP uptake (Ena1^STAA^-GFP; Fig. 6D).

As a whole, these results strongly suggest that the ST-motif is a target for phosphorylation, and that in terms of function, S^1076^ and T^1077^ were the most relevant residues.

#### Ena1 internalization depends on Yck1/2

In order to gain insight as to what kinases target the ST-patch for phosphorylation, we monitored Ena1 distribution in seven strains bearing deletions in genes encoding for different S/T kinases (*yck1-Δ1::ura3, yck2-2ts; ypk1Δ*; *ypk2Δ*; *pkh1Δ*; *pkh2Δ*; *ptk2Δ* and *kkq8Δ*). Specifically, we selected strains deficient in S/T kinases implicated in salt homeostasis and/or endocytosis. While most of them did not lead to Ena1 uptake abnormalities, a yeast casein kinase-1 (Yck1) K.O., Yck2 temperature-sensitive double mutant strain (*yck1-Δ1::ura3, yck2-2ts*; hereafter referred to as *yck^ts^*) was significantly affected for Ena1 internalization at restrictive temperature of 37⁰C (Fig. 7A). Although this result suggest that Yck function is required for the transporter uptake, It should be noted that *given the widespread function of Yck1/2 in endocytosis* (54–58)*, yck^ts^ cells show general endocytosis defects and an overall decreased fitness (even at permissive temperature).* Importantly, we reasoned that if the Ycks are directly involved in Ena1’s ST-patch phosphorylation and this is required for uptake, the Ena1^S1076D, T1077D^ (Ena1^DDST^) mutant that emulates constitutive phosphorylation of part of the ST-patch, should be able to bypass the requirement for Yck1/2-mediated phosphorylation. Indeed, in contrast to Ena1^WT^, Ena1^DDST^ was able to significantly internalize in *yck^ts^* cells even at the restrictive temperature (Fig. 7B). Further, *in vitro* phosphorylation assays showed that His6-tagged Yck1/2 proteins (purified from yeast lysates) were capable of phosphorylating the bacterially-produced, purified Ena1 ^K1090R^-Ub CT (where CT refers to <u>C</u>-<u>T</u>erminal fragment, *i.e.,* Ena1^1076-1091^; Fig. 7C).

**Figure 7:**
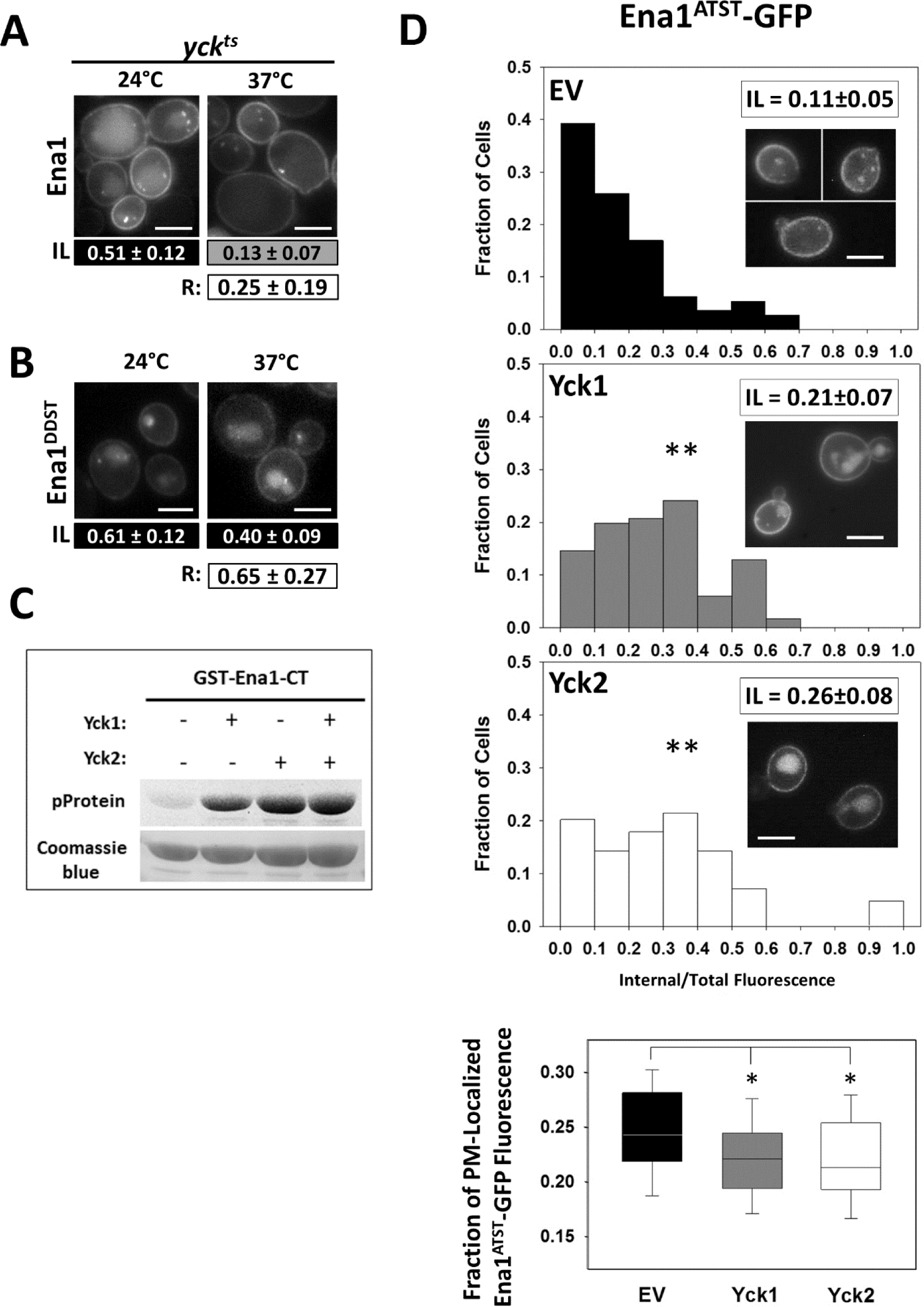
Ena1 internalization depends on the yeast casein kinases. Intracellular localization of Ena1-GFP (**A**) and Ena1^DDST^-GFP (**B**) was analyzed in *yck^ts^* cells at permissive (24°C) and restrictive (37°C) temperatures. IL indexes were estimated and analyzed as described in Fig. 1. Representative images are included. Scale bar: 5μm. “R” values in white boxes represent the ratio between the corresponding ILs for the monitored Ena1 protein at 37°C/24°C. (p<0.05). **C**. *in vitro* phosphorylation assays were performed by incubating GST-Ena1-CT protein with or without yeast purified Yck1/2 plus ATP. Phosphorylated Ena1-CT and total protein loading is shown (see *Suppl. Methods* for details)**. D**. Yeast cells expressing the Ena1^S1076A^-GFP (Ena1^ATST^) mutant were transformed with empty vector (EV) or plasmids overexpressing Yck1 or Yck2. Ena1^ATST^ relative PM accumulation and IL indexes values (and histograms) in the presence and absence of Yck overexpression, were estimated and statistically analyzed as in Fig. 1. *: p< αC=0.025— Wilcoxon’s test (Ena1^ATST^ PM-accumulation box-plots). **: p<< αC=0.025—Kolmogorov-Smirnoff test (Ena1^ATST^-GFP intracellular distribution histograms). Representative images are included. Scale bar: 5μm.

It should be noted that as a whole, the Ena1’s ST-patch is flanked by the putative casein kinase phosphorylation sites E^1073^AY<u>S</u>^1076^ and <u>T</u>^1083^E. Indeed, both S^1076^ and T^1083^ were found to be phosphorylated in whole-cell lysates of ΔΔcells expressing Ena1-GFP and Yck1/2 by mass spectrometry (Supplemental Table I). These phosphorylated S/T residues are expected to yield additional casein kinase phosphorylation sites, for example: pS^1076^TS^1078^ and T^1080^IHpT^1083^, (where pS or pT represent phosphorylated Ser or Thr, respectively) which could further propagate the post-translational modification within the ST-patch. These sites were also found phosphorylated using mass spectrometry (Supplemental Table I). This propagation mechanism is expected to continue further and to yield a highly phosphorylated ST-patch.

To further confirm the relevance of casein kinases for the internalization of Ena1, we also tested the effect of Yck1/2 overexpression on the uptake of an endocytosis-defective mutant (Fig. 7D,E). We reasoned that if Yck enzymes were involved in inducing Ena1 internalization, then their overexpression should enhance the transporter uptake. Since endocytosis of WT Ena1-GFP is highly efficient (IL≥0.8; P/T∼0.13) with a small window for improvement, we selected a transporter mutant with IL<0.5 and P/T>0.13 to improve the experiment sensitivity. Specifically, we chose the partial phosphorylation S^1076^A Ena1 mutant (Ena1^ATST^), which should be less efficient at initiating Yck-sequential phosphorylation of the ST-patch. We predicted that Yck overexpression would ameliorate such phosphorylation impairment, improving the Ena1 mutant uptake. Indeed, overexpression of either Yck1 or Yck2 led to an increase in the proportion of cells with substantial intracellular Ena1^ATST^-GFP and decreased the median accumulation of the mutant at the plasma membrane (Fig. 7D,E).

In summary our results indicate that in addition to Ub modification/recognition-deficiencies, *Ena1 internalization was also impaired*:

-by mutation of the S^1076^TST^1079^ patch to Gly or Ala (but this defect was reduced by introduction of D/E residues in positions 1076 and 1077). This ST-patch is flanked by putative casein kinase phosphorylation sites and phosphorylation of this region was verified by mass spectrometry of yeast lysates. Interestingly, overexpression of Yck1/2 improved the uptake of an Ena1 ST-patch partial mutant (S1076A).

-by Yck1/2 lack-of-function (but this defect could by bypassed by emulation of constitutive phosphorylation using the Ena1^S1076D, T1077D^ mutant). Further, purified Yck1 or 2 from yeast lysates was able to phosphorylate the Ena1 C-terminus.

Therefore, we propose that Yck-mediated phosphorylation of the S^1076^TST^1079^ patch is also required for Ena1 internalization.

### K^1090^ and the ST patch are two independent and spatially-arranged elements required for Ena1 internalization

S/T phosphorylation has been shown to be a prerequisite for ubiquitination during internalization of epsin-*independent* cargoes such as Ste6 (59). We reasoned that if the ST motif was required for Ena1 ubiquitination, then Ena1^AAST^ mutant with an *Ub fused in frame* should bypass this requirement and undergo normal internalization. However, the resulting Ena1^AAST^-Ub construction exhibited a substantial defect in uptake as compared to WT (Fig. 8A). A complementary experiment showed that Ena1^DDST, K1090R^ was not internalized efficiently either (Fig. 8A).

**Figure 8:**
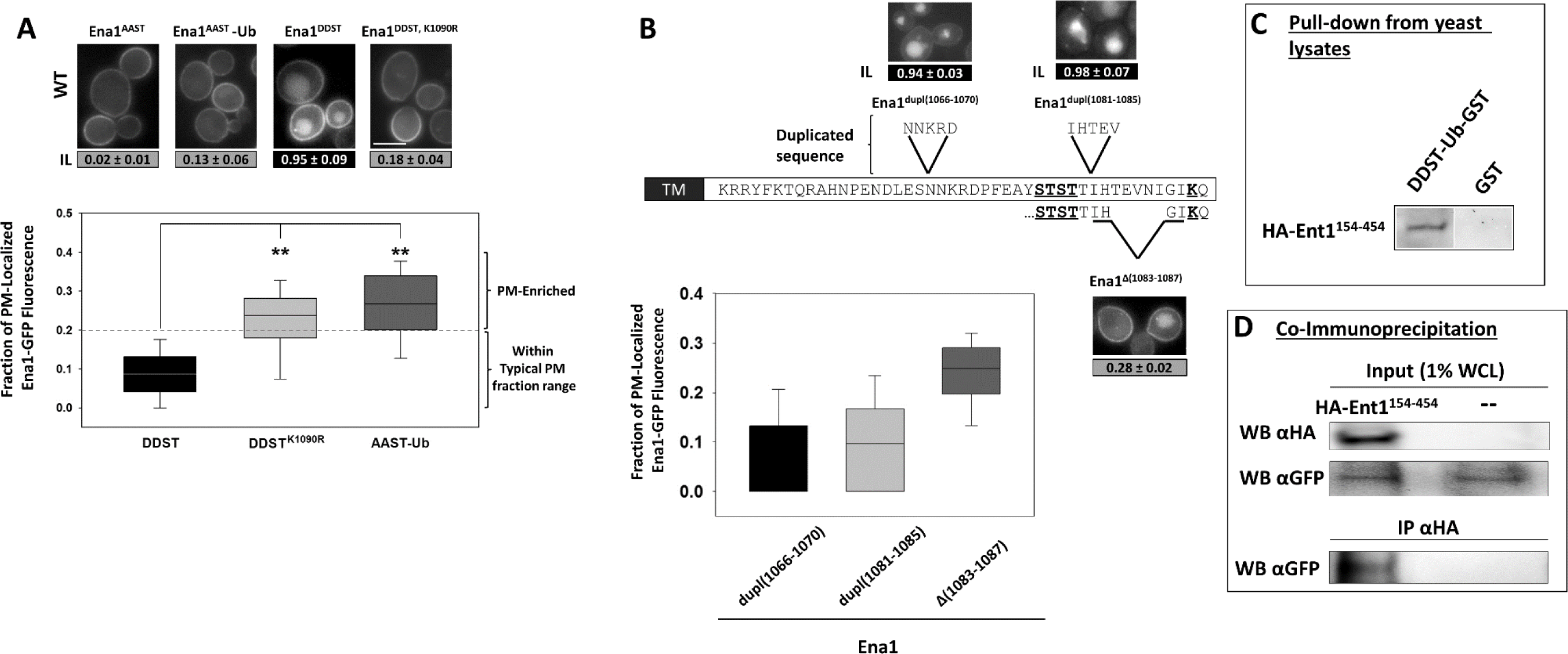
Characterization of the Ena1 STK epsin-recognition motif. The Ena1-GFP mutants indicated in **A** and **B** were expressed in yeast cells and IL indexes and relative PM accumulation values were estimated as in Fig. 1. Representative images are included. Scale bar: 5μm. Differences between the Ena1^DDST^ variant and other indicated mutants were statistically analyzed as described in *Materials and methods* and in Fig. 1 using an αC=0.01. **C**. Pull-down from yeast lysates: GSH-beads bearing immobilized GST or DDST-Ub-GST were incubated with lysates from cells expressing HA-Ent1^149-454^ as indicated in *Supplemental Methods*. The presence of bound HA-Ent1^149-454^ on beads was investigated by Western blotting with an anti-HA antibody. **D**. Co-Immunoprecipitation: These experiments were performed as described in Supplemental Materials and methods. Western blotting (WB) with the indicated antibodies of an input sample (1% of whole cell lysate used for incubation with beads) show the presence (or absence) of HA-Ent1^149-454^ and Ena1-GFP. Following immunoprecipitation with an anti-HA antibody, the presence of bound Ena1-GFP to the beads was investigated using an anti-GFP antibody (see text for details).

Although we cannot completely rule out the possibility that the ST-patch is required for ubiquitination *and* for a Ub-independent activity. Taken together these results strongly suggest that the ST-motif and the K^1090^ residue are independently required for epsin-mediated Ena1 internalization. It should be noted that these experiments do not rule out the possibility that phosphorylation of *other* determinants different from the S^1076^TST^1079^ may play a role in promoting K^1090^ ubiquitination. An important corollary conclusion is that ubiquitination is necessary but not sufficient for Ena1 internalization.

To further define characteristics and requirements of this emerging “STK” epsin-specific combined motif, we investigated if the spacing between its constitutive elements impact its function. Therefore, we modified the residue-distance between the transmembrane domain (TM) and the ST patch, as well as the residue separation between the ST patch and K^1090^ by deleting or duplicating 5-residue long segments of the Ena1 C-terminal sequence. We adopted this approach instead of using artificial spacers to minimize the possibility of introducing changes in flexibility or hydrophobicity that could independently affect Ena1 behavior. However, we cannot completely rule out the possibility that besides spacing, our results may also reflect sequence-specific effects. In fact, our data indicate that eliminating residues between the TM and the ST-patch led to ER-retention, further supporting the idea that the 1050-1070 region integrity is crucial for Ena1 proper folding. In contrast, duplication of the 1066-1070 sequence did not affect the ability of Ena1 to exit the ER and to be internalized (Fig. 8B). When the ST-K residue distance was shortened from 10 to 5 (Δ1083-1087) led to deficiencies in Ena1 uptake, whereas duplication of a 5 residue-sequence (1083–1087) did not affect internalization (Fig. 8B). These results suggest that a TM-ST spacing ≥29 and ST-K ≥10 sustained proper function of the STK sequence as a combined, epsin-specific endocytic motif.

Interestingly, known epsin mammalian cargoes such as VEGFR2, Delta-like 1 and ENaC bear C-terminal, cytoplasmic sequences with similar composition (rich in Ser, Thr *or acidic residues*) at the appropriate distance from a lysine, constituting potentially analogous STK motifs to the one identified in Ena1 (Supplemental Fig. 4). In fact, the “ST”-like motifs from these mammalian proteins were able to functionally replace Ena1 ST-patch sustaining epsin-dependent Ena1 internalization (Supplemental Fig. 4).

Finally, and to evaluate if epsin was able to associate with a transporter mutant emulating the double post-translationally modified STK Ena1 motif, we performed pull-down and co-immunoprecipitation (co-IP) experiments.

#### Epsin pull-down experiments from yeast whole cell lysates (WCL)

Glutathione-beads bearing immobilized GST or GST-Ub-Ena1^S1076D, S1077D^CT as described above and incubated with WCL from cells expressing HA-Ent1^149-454^ (this truncation was preferred to the more aggregation-prone, ENTH-domain containing, full-length Ent1; see Supplemental Materials). After incubation and washes, the beads were boiled in sample buffer, resolved by SDS-PAGE and the presence of bound HA-Ent1^149-454^ investigated by Western blotting with anti-HA antibodies. Results shown in Figure 8C indicate that recombinant cytoplasmic C-terminal tail of Ena1 “constitutively phosphorylated/ubiquitinated” was able to interact with *in vivo*-produced epsin protein from cell lysates

#### Co-immunoprecipitation

Although co-IP of intrinsically low affinity, sorting machinery-cargo complexes is challenging (and further aggravated by the presence of 10 transmembrane domains in Ena1) we were able to co-immunoprecipitate full-length Ena1^S1076D, S1077D^-Ub – GFP with HA-tagged Ent1^149-454^ using an anti-HA antibody (a representative experiment is shown in Fig. 8D). This result indicated that epsin was capable of *in vivo* recognition of constructions that emulate constitutively phosphorylated/ubiquitinated STK from Ena1.

## DISCUSSION

Epsin is an endocytic protein essential for cell viability in yeast and required for embryo development in metazoans (60). The function of this protein family as endocytic adaptor is necessary for activation of the Notch signaling pathway (24, 26, 27) and for the temporal-spatial coordination of endocytosis and RhoGTPase signaling (61–63). Epsin endocytic function is required for VEGFR2 (23), VEGFR3 (64) and ENaC internalization (28, 29) and (although not essential) is known to contribute to the uptake and regulation of EGFR, insulin receptor, mu opioid receptors, dopamine transporter, protease activated receptor1, Ste2 and others (65–74)

However, how epsin (or other Ub-binding endocytic adaptors) recognize their specific cargoes among multiple ubiquitinated proteins is unknown. Here, we demonstrate the existence of a coincidence detection mechanism that assures specific recognition of ubiquitinated cargo by yeast epsins. We speculate that other adaptors, such as the Eps15-homolog Ede1, require post-translational modification of other determinants (in addition to ubiquitination) for recognition of their corresponding specific cargoes.

In fact, this study unveils several important aspects of yeast epsin-mediated internalization of cargo. We have established the Na^+^ pump Ena1 as the first epsin-specific cargo identified in *Saccharomyces cerevisiae* and showed that epsins mediate internalization of Ena1-GFP upon salt-stress relief. Specific residues within the Ena1 C-terminus (S^1076^TST^1079^ and K^1090^) critical for its epsin-mediated internalization were also mapped. Furthermore, our data strongly suggests that the K^1090^ residue is a target for Art3/Rsp5-mediated ubiquitination while the ST-patch is a target for yeast casein kinase (Yck1/2)-mediated phosphorylation. Our results also suggest that Ena1 S^1076^-T^1079^ phosphorylation was not a prerequisite for K^1090^ ubiquitination, but instead that both determinants are independently required for epsin-mediated Ena1 internalization.

Therefore, and very importantly, our data indicate that ubiquitin is not sufficient for Ena1 uptake and suggest the existence of a coincidence detection mechanism for recognition of ubiquitinated cargoes by epsin.

Our results show for the first time that ubiquitination-independent determinants present in the cargo (in this case, the ST-motif of Ena1) are required for Ub-binding adaptor (*e.g.,* epsin) specificity.

Indeed, our results indicate that internalization of Ena1 by epsins does not rely on direct recognition of sequence-encoded sorting signals (such as Y-or L-motifs (75)), but on a double post-translational modification (ubiquitination and phosphorylation) of specific cargo regions.

On the one hand, Ena1 internalization was dependent on the presence of the arrestin-related adaptor Art3 linking the yeast ubiquitin ligase Rsp5 to this process. This is in agreement with the well-known role of Rsp5 (the only HECT Ub-ligase in *S. cerevisiae)* in endocytosis (76). Although Art9 has been shown to be important for the regulation of Ena1 (53), our data indicates that is not crucial for internalization of the transporter. We speculate that Art9 will be involved in the ubiquitination of K residues relevant to other steps of intracellular traffic while Ena1 K^1090^-Ub is required for removal from the plasma membrane. Indeed, Ub-fusion in frame of an Ena1 K1090R mutant re-established cargo uptake from the cell cortex, but did not induced direct Golgi-to-MVB/vacuole traffic or PM recycling.

On the other hand, our results showed that Ena1 S^1076^TST^1079^ patch can be indeed phosphorylated *in vivo* and strongly suggested that yeast casein kinases are involved in the transporter uptake. Although we cannot completely rule out involvement of other serine/threonine kinases in Ena1 internalization, our findings are in agreement with the well-known role of Ycks in endocytosis (54–58).

It has been clearly shown that Yck activity is a pre-requisite for ubiquitination of the epsin-independent cargo Ste3 (54, 77); however, our data indicated that Ena1-ubiquitination does not require ST-patch phosphorylation. Nevertheless, *it is possible that phosphorylation of other determinants in Ena1 cytoplasmatic domains could support K^1090^ ubiquitination*.

Therefore, it is tempting to speculate that independence between phosphorylation of this specific ST patch and ubiquitination in Ena1 is a landmark of epsin-specific internalization.

Our results also suggest the existence of an epsin-recognized, combined STK motif with specific required spacing between its components and from the last transmembrane domain. We speculate, that such requirements may reflect steric/spatial constraints for binding the transmembrane, STK-bearing cargo by the membrane-bound epsins.

Searching the *Saccharomyces cerevisiae* database (http://www.yeastgenome.org/) we found several transmembrane proteins displaying putative STK-like sequences with similar spacing and position than Ena1 (including the whole Ena protein family). However, the functionality of putative STK motifs for epsin-mediated internalization must be experimentally tested (and it will be the focus of future investigations) as unknown context factors and additional determinants might also play an important role in the control of plasma membrane cargo levels.

Indeed, the complexity of the trafficking information packed in the cytosolic (and possibly transmembrane) portions of membrane proteins should not be underestimated. Important work done in the Chen lab clearly indicated that in mammalian epsins specificity determinants for the recognition of the receptor tyrosine kinase VEGFR2 are embedded within the UIM sequence environment (78–80). Further, it should be kept in mind that multi-spanning proteins like Ena1 usually contain additional/critical determinants in their intracellular loops; these regions may play a role in the recruitment of post-translational modifying enzymes; *e.g.,* kinases, ubiquitin ligases/ARTs; or proteins that may assist in cargo recognition. A binding site for the fly Ub-ligase *Mind bomb* has been found within the cytosolic domain of the epsin cargo protein Delta (81); very interestingly, this region is flanked by ST/K-like elements. Further, the simple transplantation of the Ena1 STK motif was not enough to convert Ste3 into an epsin-specific cargo but rendered a plasma membrane-localized hybrid protein. A similar result was observed upon replacement of Ena1 C-terminus for the analogous Ste3 sequence (data not shown). These observations further suggest a complex organization of partially overlapping motifs required for cargo internalization.

In terms of the specific regulation of the plasma membrane levels of the Na^+^ pump Ena1, we propose the following working model (Fig. 9): Upon salt stress Ena1 is expressed and translocated at the cell cortex to contribute to the efflux of sodium (43) (Fig. 9). When salt levels decrease, Ena1 is post-translationally modified i) by ubiquitination at K^1090^ (by an Art3/Rsp5 complex) and ii) by phosphorylation at the S^1076^TSTT^1080^ patch (by casein kinases) (Fig. 9). The double modified C-terminus of Ena1 then becomes competent for specific recognition of yeast epsins and retention at nascent endocytic sites by these Ub-binding adaptors. Epsins, in turn, contribute to the recruitment, or further binding, of accessory factors and the clathrin coat (60), which eventually leads to the pinching-off of endocytic vesicles thereby removing Ena1 from the plasma membrane (Fig. 9).

**Figure 9:**
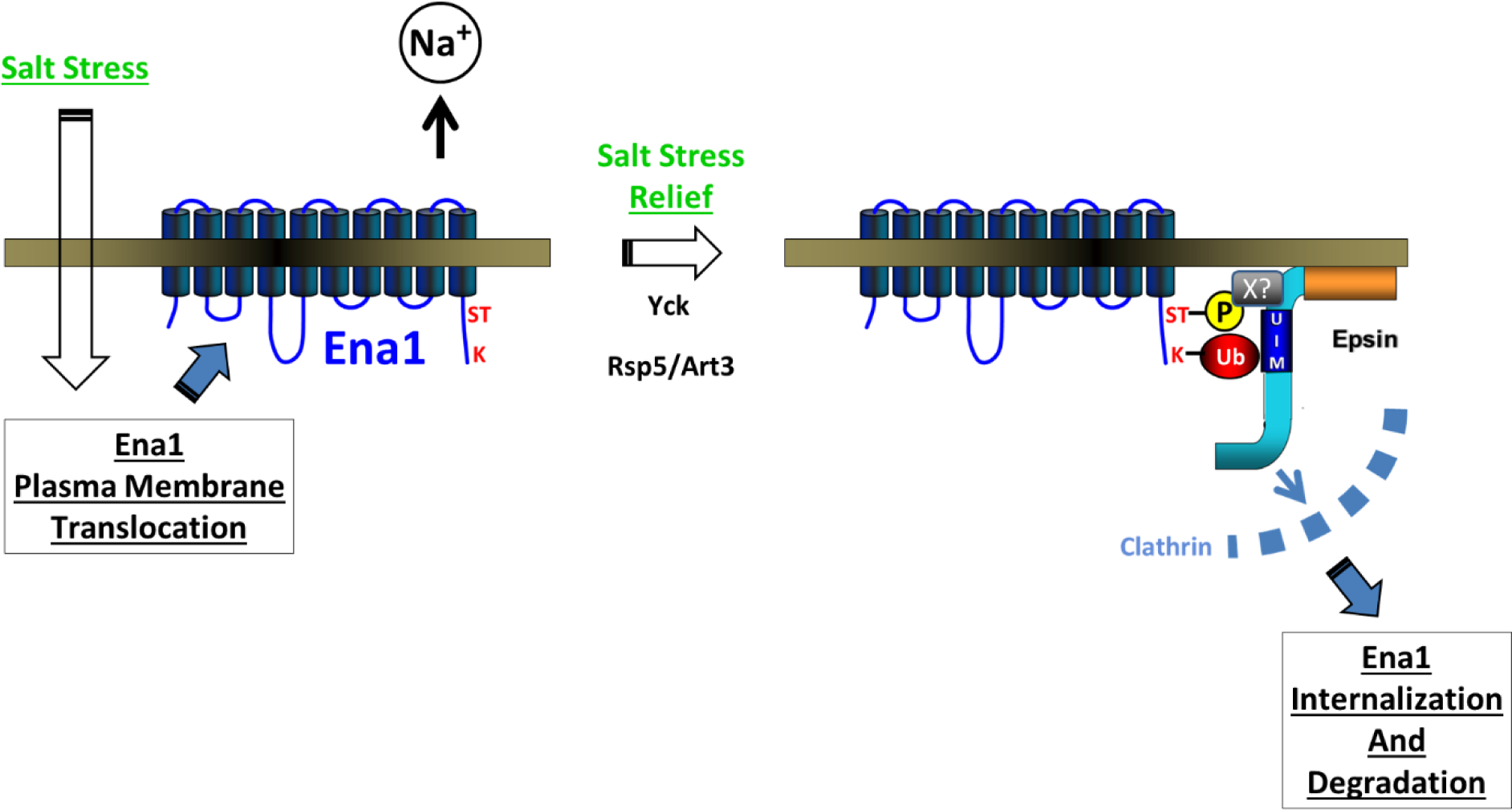
Working model for Epsin-dependent Ena1 internalization. Cartoon depicts the proposed mechanism that assures specific recognition of the ubiquitinated cargo Ena1 by Ub-binding adaptor epsin (see text for details).

We believe that this model, supporting findings and reagents will be the basis of future investigations that will further our understanding of endocytic function of epsin in yeast.

Importantly, one could speculate that alternative coincidence detection mechanisms confer recognition specificity by other ubiquitinated-cargo/ubiquitin-binding adaptor combinations (*e.g.,* involving Ede1). Finally, since mammalian STK motifs sustained Ena1 internalization in yeast, we also speculate that this strategy might be conserved across evolution. Nevertheless, these emerging hypotheses will need to be tested taking into account complexity (see above) and potential redundancy of endocytic signals and endocytic adaptors.

## ACKNOWLEDGEMENTS

We thank Drs. Chris Staiger, Donald Ready, Henry Chang and Tony Hazbun (Purdue University) for stimulating discussions and/or critical reading of the manuscript. We also thank Drs. Jeremy Thorner (UC Berkeley), Lucy Robinson (Louisiana State University), Ingrid Wadskog (Göteborg University) and Martha Cyert (Stanford) for reagents. This work was supported by the National Science Foundation under Grant No. 1021377 to RCA and by the Center for Science of Information (CSoI), an NSF ST Center, under grant CCF-0939370.

## MATERIALS & METHODS

### Reagents and DNA constructs

Materials were purchased from Fisher Scientific (Fairlawn, NJ) or Sigma (St. Louis, MO) unless stated otherwise. Plasmids and strains used in this study are listed in supplemental tables II and III. DNA constructs were prepared using standard techniques. Site directed mutagenesis was done using a QuikChange^®^ Lightning Site Directed Mutagenesis kit (Agilent Technologies, Inc., Santa Clara, CA).

### Yeast culture conditions and transformation procedures

Yeast SEY6210-derived (MATα *leu2-3,112 ura3-52 his3-Δ200 trp1-Δ901 suc2-Δ9 lys2-801; GAL*) and other strains were grown overnight at 30°C (or room temperature for temperature-sensitive strains) with shaking at 250 RPM in standard yeast extract–peptone–dextrose (YPD) or synthetic selective (amino acid dropout) media supplemented with dextrose and lacking appropriate amino acids for plasmid maintenance. Yeast was transformed by the Li-Acetate method following standard techniques.

### Microscopy

Images were acquired using a Zeiss Axiovert 200M microscope equipped with Zeiss Axiocam MRm monochrome digital camera and Carl Zeiss Axiovision image acquisition software (version 4.4). For imaging *MET25* promoter driven Ena1-GFP derived constructs, cells were grown overnight in selective media in the presence of 2mM methionine. The cultures were diluted, washed with sterile H2O and Ena1 expression was induced by growing the cells in selective media without methionine for 5h. A cell culture volume containing the equivalent of 2-5 OD600nm of cells expressing fluorescently-tagged proteins were pelleted and resuspended in 50-100μl selective media, 10μl of cell suspension was spotted on a pre-cleaned slide, covered with a 22X22mm coverslip and imaged with appropriate filters as ∼0.3μm-spaced Z-stacks.

### Statistical analysis

Was performed as described in Suppl. methods and in (82).

## Supplemental Information

### Supplemental Material and Methods

#### FM4-64 staining

To visualize the endocytic pathway and vacuolar membrane, staining using FM4-64 was performed according to (1). Briefly, cells were grown overnight to 0.3-0.8 OD/ml in selective media. 1 mL of cells were pelleted and chilled on ice, followed by the addition of cold FM4-64 (1:40 dilution from a stock of 1mg/ml in DMSO) in a final volume of 50μl of selective media. The cells were incubated for 15-20min on ice and subsequently imaged (t = 0min). Cells were supplemented with 900μl of fresh pre-warmed selective media and allowed to incubate at 30°C for 30min. At this time point, cells were quickly chilled on ice prior to imaging at 100X using GFP and rhodamine filters.

#### Quantitative analysis of Ena1-GFP localization

To quantify the localization of fluorescently-tagged Ena1, we used an approach based on the analysis of fluorescence signal distribution within cells in randomly acquired images. First, we manually outlined each cell and measured the total GFP-fluorescence intensity (T) using ImageJ software. T represents the sum of fluorescence contributions associated with internal compartments (I), plasma membrane-localized (P) and cytosolic background (C). The fluorescence intensity inside of the cell (IC=I+C) was measured by outlining the cell of interest excluding the peripheral region, while the P component value was estimated subtracting IC from T. The average contribution of cytosolic fluorescence was quantified by measuring and averaging the fluorescence intensity of three rectangular regions within the cytosol of each analyzed cell. This value multiplied by the cell area yielded an integrated cytosolic background intensity (C). Internal fluorescence intensity was calculated as I=IC - C. The I component arises from endocytic uptake of Ena1-GFP as shown by >95% colocalization with the hydrophobic dye FM4-64, (Supplemental Fig. 1). All measurements were performed on a single plane with the best focus of cell periphery. This analysis was performed on at least 30 cells and each experiment was performed at least 3 times.

The results were processed and represented in two complementary ways:

1. *Histograms (fraction of cells vs. I/T values)* provided a measurement population distribution. In addition to direct histogram comparisons, an ‘Intracellular Localization’ (IL) index was defined as the cumulative fraction of cells with intracellular fluorescence greater 50% of the total fluorescence; *i.e.,* fraction of cells present at the right-hand side of a line positioned at I/T=0.5 over at least 3 histograms. This value integration procedure yielded a normally distributed, very convenient *descriptor of population skewness with respect to Ena1-GFP localization*. The higher the IL, the higher *the proportion of cells* with cargo enrichment in intracellular compartments.
2. *P/T median values* provided a *measure of Ena1-GFP plasma membrane accumulation in cells*. Since P=T-IC (see above), calculation of this quantity does not require estimation of cytosolic background; *i.e.,* decreasing error propagation. Although showing typical cell-to-cell biological variation (and following a non-normal distribution), P/T values were found to be fairly independent of Ena1-GFP levels of expression (Supplemental Fig. 2). Nevertheless, highly overexpressing cells were excluded from all analyses. A P/T median value ≤0.2 (Supplemental Fig. 2) was considered within the limits for typical (normal) Ena1-GFP PM fraction range.

It should be noted that populations with similar ILs can show substantially different P/T median values.

#### Salt stress relief assay

WT and *ent1Δent2Δ* cells expressing Ena1-GFP were grown overnight in selective media supplemented with 0.2mM methionine to maintain low levels of Ena1 expression (controlled by *MET25* methionine-repressible promoter). Cells were grown to early log phase, washed and transferred to media lacking methionine for 2h, thereby inducing high Ena1 expression for a short time. Next, 0.5M NaCl was added to the cells to simulate salt stress and simultaneously supplemented with 2mM methionine to suppress the *MET25* promoter (stopping Ena1 production), cells were incubated under these conditions for 3h. This was done to ensure that the Ena1-GFP produced by the cells would be recruited and maintained at the cell membrane to pump out excess sodium. The cells were imaged (time 0), and then washed to remove NaCl excess (“salt stress relief”—under these conditions we expected Ena1 to be internalized) and resuspended in media containing 2mM methionine. Cells were imaged over time and the amount of peripherally localized Ena1 was quantified as described in Materials and methods.

#### Flow Cytometry

Wildtype or *ent1Δent2Δ* cells transformed with Ena1-GFP plasmid were cultured in synthetic complete medium lacking methionine at 30°C overnight. The cells were diluted to 0.2 OD600nm/ml with fresh media and grown for 2h. Cells were supplemented with methionine to a 2mM final concentration, and a sample was taken immediately after (“0h”), and another after 5h incubation at 30°C (“5h”). After staining with propidium iodide (BD Biosciences) for 10min, 5x10^4^ cells were analysed using a FC500 flow cytometer (Beckman-Coulter) with FL1 and FL3 detectors. The results were processed using FlowJo v7.6 software.

#### Cell lysate preparation

Cells were grown in selective media and spheroplasted following standard techniques (2). Briefly, ∼50OD600 were harvested and incubated for 10min at room temperature in softening solution (0.1M Tris pH 9.4, 10mM DTT, 5mM NEM). Cells were then resuspended in spheroplasting solution composed of YPD medium containing 1M sorbitol, 1unit of zymolyase per OD600, protease inhibitor cocktail (Pierce) and 0.1mM AEBSF (4-(2-aminoethyl)benzenesulfonyl fluoride) for 30min at 30°C. Spheroplasts were washed once with PBS containing 1M sorbitol, pelleted by centrifugation at 300xg for 5min and dounce-homogenized with in HEPES/KOAc lysis buffer (0.2M sorbitol, 50mM KOAc, 2mM EDTA, 20mM HEPES pH 6.8, 0.2% Triton X100, 50mM NEM, protease inhibitor cocktail).

#### Western blotting

##### Ena1-GFP detection by western blotting

cells were induced to express Ena1-GFP (WT or mutant) by maintaining them in medium without methionine for 5h. The equivalent to 10 OD600 of cells were collected and resuspended in 200ul of 0.2M NaOH for 15min at room temperature. Cells were spun down at 1000xg for 2min and resuspended in 50μl of 2x Laemmli buffer with 8% SDS (100mM Tris–HCl, 8% SDS, 40% glycerol, 2% mercaptoethanol, 0.005% bromophenol blue, pH6.8) and incubated at 55°C for 15min. the equivalent to 50μl of 0.15-0.25mm acid washed beads were added to the sample and vortex for 1min at room temperature. The samples were chilled on ice for 2min and spun down at 300xg for 5min. The supernatants were collected and resolved in 8% SDS-PAGE gels, transferred onto nitrocellulose followed by immuno-blotting using an anti-GFP antibody (Thermo Fisher A-11122) at a 1:1000 dilution. The loading control, VDAC1 (porin) was detected by using a specific mouse monoclonal antibody (Clone 16G9E6BC4, Thermo Fisher) at 1:1000 dilution. Blots were developed using enhanced chemiluminescence detection system.

#### Crude Membrane Preparation

Cells expressing Ena1-GFP were grown overnight in the presence of 0.2mM methionine. Cells were diluted in medium without methionine to final 0.5 OD600/ml and to induce Ena1-GFP expression for 5h. 200 OD600 of cells were collected by centrifugation at 1000g for 5min, resuspend in ice-cold water and aliquot to 10 OD600 /tube. The cells were then pelleted down at 2000xg for 2min, 0.15-0.25mm acid-washed glass beads (equivalent to a 100μl volume) and 1ml of ice-cold lysis buffer [30mM Tris-HCl (pH8.5), 5mM EDTA (pH 8.0), 25mM DTT, 250mM NaF,2X protease inhibitor, 2X phosphatase inhibitor cocktails (Pierce)] were added to each tube. The cells were broken by vortexing for 3min, then chilled on ice for 2min, repeating the cycle twice. The crude membranes were collected by pelleting the lysates at 300xg for 5min, the supernatants were pooled and centrifuged at 20,000xg 30min at 4°C. The pellet was resuspended in 1ml of lysis buffer containing 0.1% SDS.

#### Mass spectrometry

##### Protein extraction and phosphopeptide enrichment

GFP-tagged Ena1 was purified by incubating extracted crude membranes with GFP-Trap agarose beads (ChromoTek, NY) for 1h at 4°C. The GFP-Trap beads were washed with RIPA buffer [50mM Tris-HCl (pH 7.5), 150mM NaCl, 1% NP-40, 0.5mM EDTA, 0.1% SDS, 0.5% sodium deoxycholate, and 50mM NaF] three times, and enriched Ena1-GFP proteins were denatured in 8M urea and reduced with 5mM dithiothreitol for 30min at 37°C. The proteins were further alkylated in 15mM iodoacetamide for 1h in the dark at room temperature and then digested with proteomics grade trypsin at a 1:100 (w/w) ratio overnight at 37°C. The digested peptides were desalted by SDB-XC reverse phase-Stage Tips (3).

Phosphopeptides from Ena1-GFP were enriched by polyMAC (4). The eluted phosphopeptides were dried in a SpeedVac and stored at -20 °C.

##### LC-MS/MS analysis and data analysis

The phosphopeptides were dissolved in 4ul of 0.1% formic acid and injected into an Easy-nLC 1000 (Thermo Fisher Scientific). The mobile phase buffer consisted of 0.1% formic acid in ultra-pure water (Buffer A) with an eluting buffer of 0.1% formic acid in 80% ACN (Buffer B) run over a linear 90min gradient with a flow rate of 250nl/min. The Easy-nLC 1000 was coupled online with a Velos LTQ-Orbitrap mass spectrometer (Thermo Fisher Scientific). The mass spectrometer was operated in the data-dependent mode in which a full-scan MS (from m/z 300-1500 with the resolution of 60,000 at m/z 400) was followed by top 15 MS/MS scans of the most abundant ions. The raw files were searched directly against *Saccharomyces cerevisiae* database (UniprotKB) with no redundant entries using both SEQUEST HT and BYONIC algorithm on Proteome Discoverer^TM^ version 2.0 (Thermo Fisher Scientific). Peptide precursor mass tolerance was set at 10 ppm, and MS/MS tolerance was set at 0.6 Da. Search criteria included a static carbamidomethylation of cysteines (+57.0214 Da) and variable modifications of (1) oxidation (+15.9949 Da) on methionine residues, (2) Gln to pyro-Glu (-17.027 Da) at N-terminus of peptide, (3) acetylation (+42.011 Da) at N-terminus of protein, and (4) phosphorylation (+79.996 Da) on serine, threonine or tyrosine residues were searched.

Search was performed with full tryptic digestion and allowed a maximum of two missed cleavages on the peptides analyzed from the sequence database. Relaxed and strict false discovery rates (FDR) were set for 0.05 and 0.01, respectively.

#### Recombinant protein purification

Bacterially produced recombinant proteins were purified based on the procedure described in (5). Rosetta cells (Novagen) were transformed with GST- or His6-tagged constructs (listed in supplemental table I) and protein production was induced by incubation with 0.2mM (final concentration) isopropyl-β-D-thiogalactopyranoside (IPTG) for 5h at 30⁰C. The cells were harvested and resuspended in PBS containing 0.1% Tween (PBST) plus 5% glycerol, protease inhibitor cocktail and 0.1mM AEBSF. Cell lysis was induced by incubating the cell suspension with 1mg/ml lysozyme (4⁰C, 15min) followed by sonication. Clarified lysates were incubated either with Ni-NTA beads in the presence of 5mM imidazole (for purification of His6-tagged proteins) or with glutathione coupled beads (for purification of GST-tagged proteins), for 2h at 4⁰C. The beads were washed 4x with PBST + imidazole (in increasing concentrations of 5mM, 10mM, 15mM and 20mM) for His6-tagged proteins, and 4x with PBST alone for GST-tagged proteins. Elution was performed with either 500mM Imidazole (His6-tagged proteins) or 50mM glutathione (GST-tagged proteins). The supernatant containing the purified tagged proteins were collected, and the imidazole from His6-tagged, and glutathione from GST tagged protein fractions were removed using Zeba^TM^ spin desalting columns (Thermo Scientific, Waltham, MA).

##### in vitro kinase assay

C-terminal His6-tagged Yck1/2 proteins were expressed in yeast and purified from whole cell lysates (equivalent to ∼100 OD600 of cells) using Ni^2+^-NTA beads and eluted with 500μl of 500mM imidazole and desalted into 500μl of 2x kinase buffer [100mM Tris-HCl pH 7.5, 20mM MgCl2, 2mM DTT]. Purified Yck1/2 proteins were incubated with GST-Ena1^K1090R^CT-Ub immobilized on 50μl of glutathione beads at 30°C for 45min in 1x kinase buffer with 5mM ATP. The beads were then washed 3 times with 1x kinase buffer and boiled with 25μl of 2x protein sample buffer. Samples were resolved with SDS-PAGE and phosphorylated proteins were detected using Pro-Q® Diamond Phosphoprotein Gel Stain kit (ThermoScientific, Waltham, MA) following manufacturer’s instructions.

#### Pull-down assay with cell lysate and recombinant proteins

Overexpression of HA-Ent1^149-454^ under the regulation of a *MET25* promoter was induced by growing the cells overnight in selective media in the absence of methionine. For these and for co-immunoprecipitation experiments, the Ent1 truncation indicated above was preferred to the more aggregation-prone, ENTH-domain containing, full-length Ent1. Cell lysates were prepared as described above. The samples were centrifuged at 8000 rpm for 5min, and clarified lysates were incubated with GST or GST fusion proteins immobilized on glutathione coupled beads for 2h at room temperature. The beads were washed 5 times with PBST, resuspended in Laemmli’s protein sample buffer, followed by SDS-PAGE resolution and immunoblotting. A mouse monoclonal α-HA antibody was used (Covance, Princeton, NJ) at 1:2500 dilution.

#### Co-immunoprecipitation assay

Yeast cells expressing *MET25* promoter driven Ena1-GFP transformed with empty vector or a plasmid encoding for HA-tagged Ent1^149-454^ (under endogenous promoter), were grown overnight in selective media in the absence of methionine. Cells were lysed by spheroplasting and dounce homogenization as described above and subjected to cross-linking using 2mM DSP [Dithiobis (succinimidyl propionate)] for 30min at RT with constant rotation. The crosslinking reaction was quenched by incubating with 50mM Tris-HCl pH 7.5 for 15min. For protein extraction mild alkali treatment was used (6). NaOH was added to the mix to a final concentration of 0.2M and incubated for 5min at RT. Samples were centrifuged at 8000rpm for 5min and the supernatant was discarded. The pellet was resuspended in extraction buffer (0.06M Tris-Hcl pH7.5, 1% SDS, 5% glycerol, 2% β-mercaptoethanol, supplemented with N-ethylmaleimide, protease and phosphatase inhibitors) and boiled for 3min. Following centrifugation at 8000rpm for 5min, the clarified supernatant was diluted at least 20 times in 1M Tris-HCl pH 7.5. α-HA coupled beads (EZview red anti-HA affinity gel, Sigma) were used to immunoprecipitate HA-tagged proteins (at 4⁰C for 2h). Following 3 washes, the beads were boiled in Laemmli buffer for 5min. Samples were resolved by SDS-PAGE and immunoblotted. A rabbit monoclonal α-GFP antibody (Invitrogen, Carlsbad, CA) at 1:10 dilution was used to detect GFP-tagged proteins from the immunoprecipitates. HA tagged proteins were detected using a mouse monoclonal α-HA antibody (Covance, Princeton, NJ) at 1:2500 dilution.

#### Statistical analysis

When appropriate, the magnitude of errors associated with values derived from algebraic operations using experimentally measured quantities were calculated following standard rules of error propagation. Statistical significance of differences between fluorescence distribution histograms were analyzed using the Kolmogorov-Smirnoff test. The student’s t-test was used to evaluate differences among normally distributed ILs, while the Wilcoxon’s test was used to evaluate the significance between non-normal P/T of value samples. Bonferroni’s correction for multiple comparisons was performed whenever applicable [αC=p/n; n being the number of comparisons].

## Legends to Supplemental Figures

**Supplemental Fig.1.**
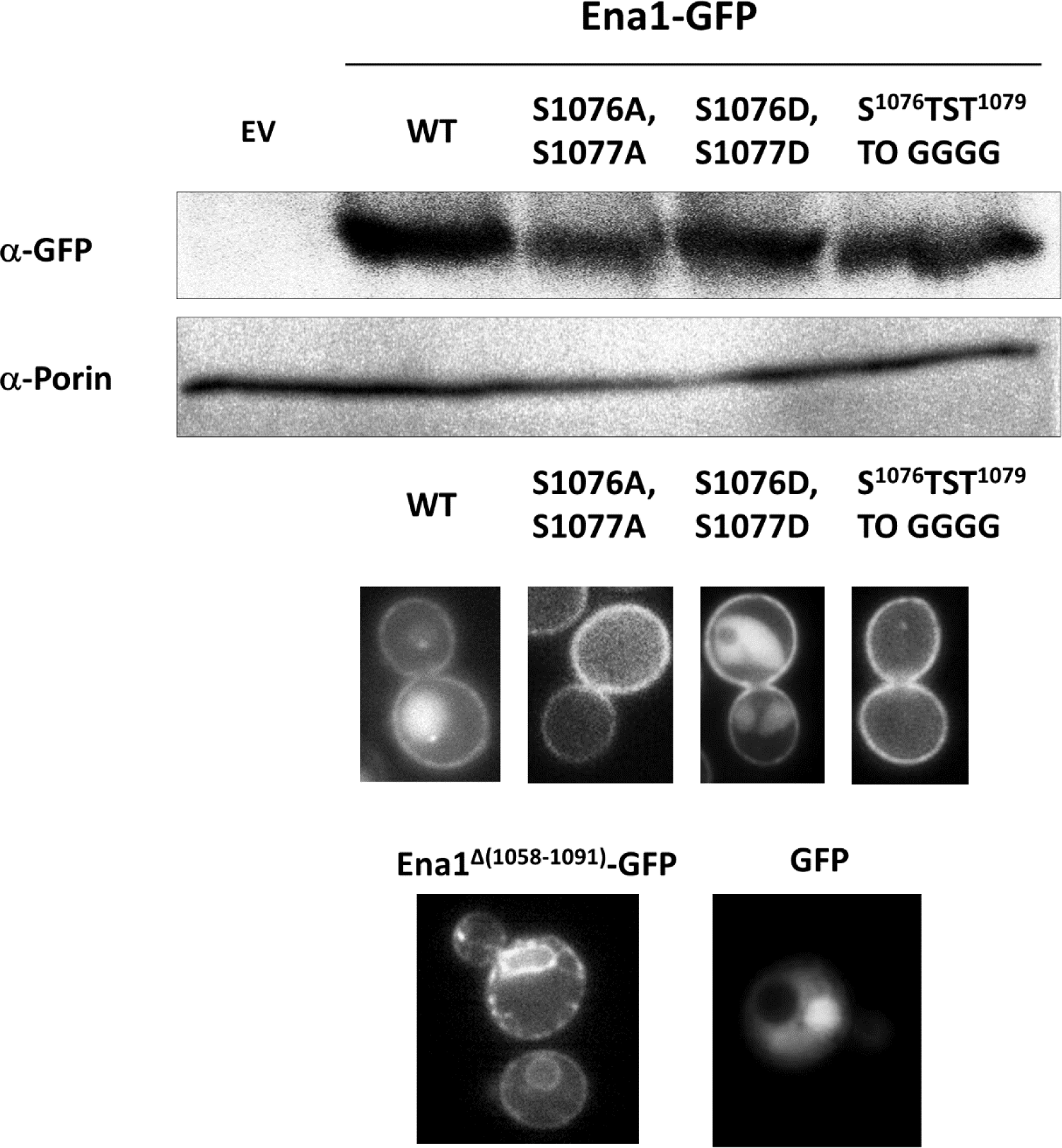
The stability and expression levels of Ena1-GFP WT or mutants were monitored by Western blotting (upper panel) and the presence/analysis of their intracellular distribution patterns by microscopy (lower panels). Western blotting was conducted as described in *Supplemental materials and methods* on whole cell lysates from cells transformed with empty vector (EV) and plasmids encoding for Ena1-GFP WT and mutants (only 3 are shown as example). Presence of Ena1-GFP was investigated using an anti-GFP antibody, while the signal from the mitochondrial protein porin was used as loading control. In addition to protein levels, the different Ena1-GFP variants were analyzed/evaluated by their intracellular distribution. Specifically, to be considered for this study, the evaluated proteins were also expected to show a pattern substantially different from plain GFP or typical ER-retention pattern (*e.g.,* Ena1^Δ(1058-1091)^.

**Supplemental Fig.2.**
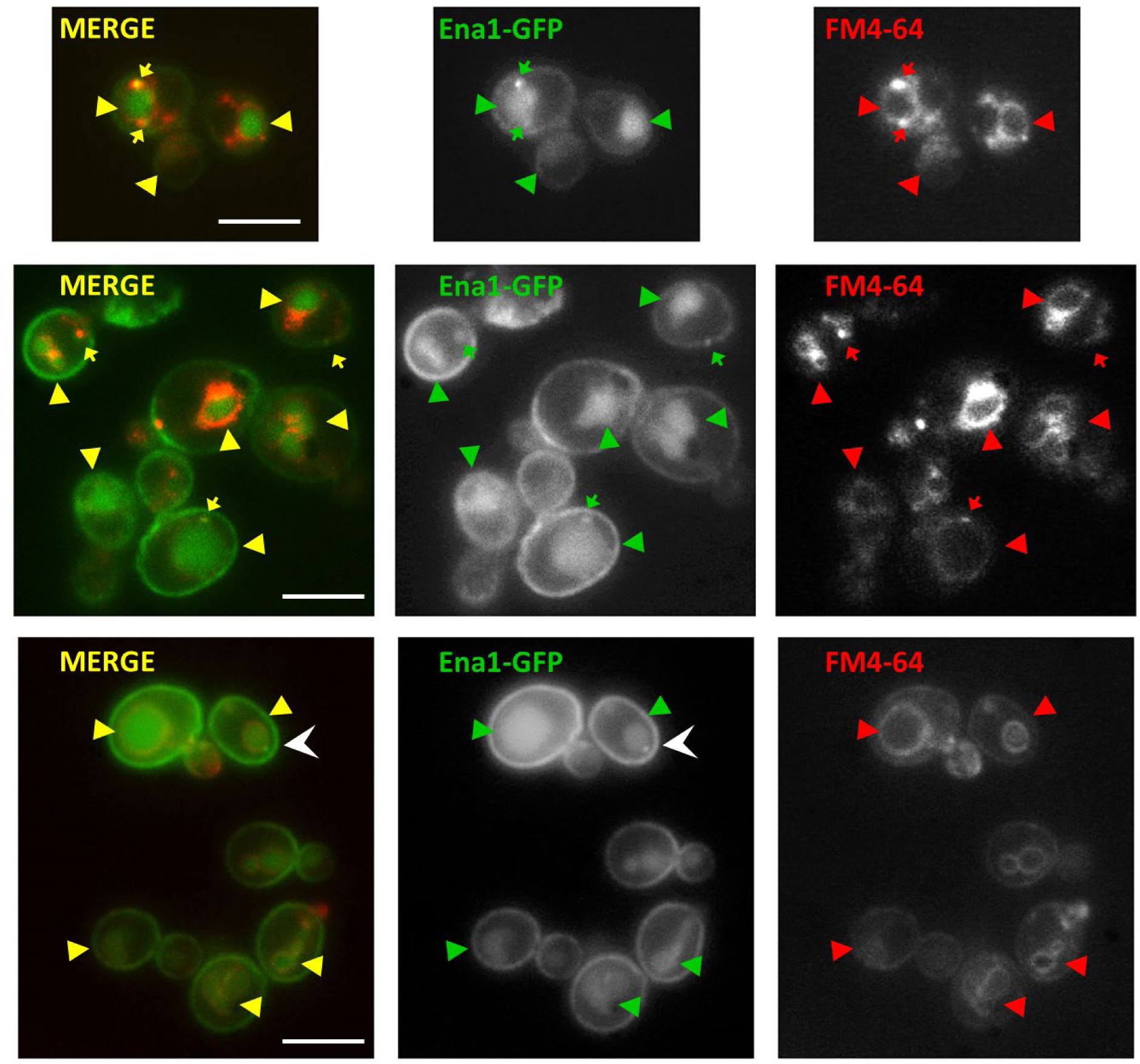
Cells expressing Ena1-GFP displayed intracellular structures labeled with the lipophilic dye FM4-64 used to mark the endocytic pathway. These Ena1-GFP positive compartments colocalized with FM4-64 (95%) in vacuoles (arrowheads) and punctate structures (arrows). White arrowhead points to one example of the very few Ena1-GFP compartments that do not colocalize with FM4-64; these intracellular structures may represent a secretory compartment.

**Supplemental Fig.3.**
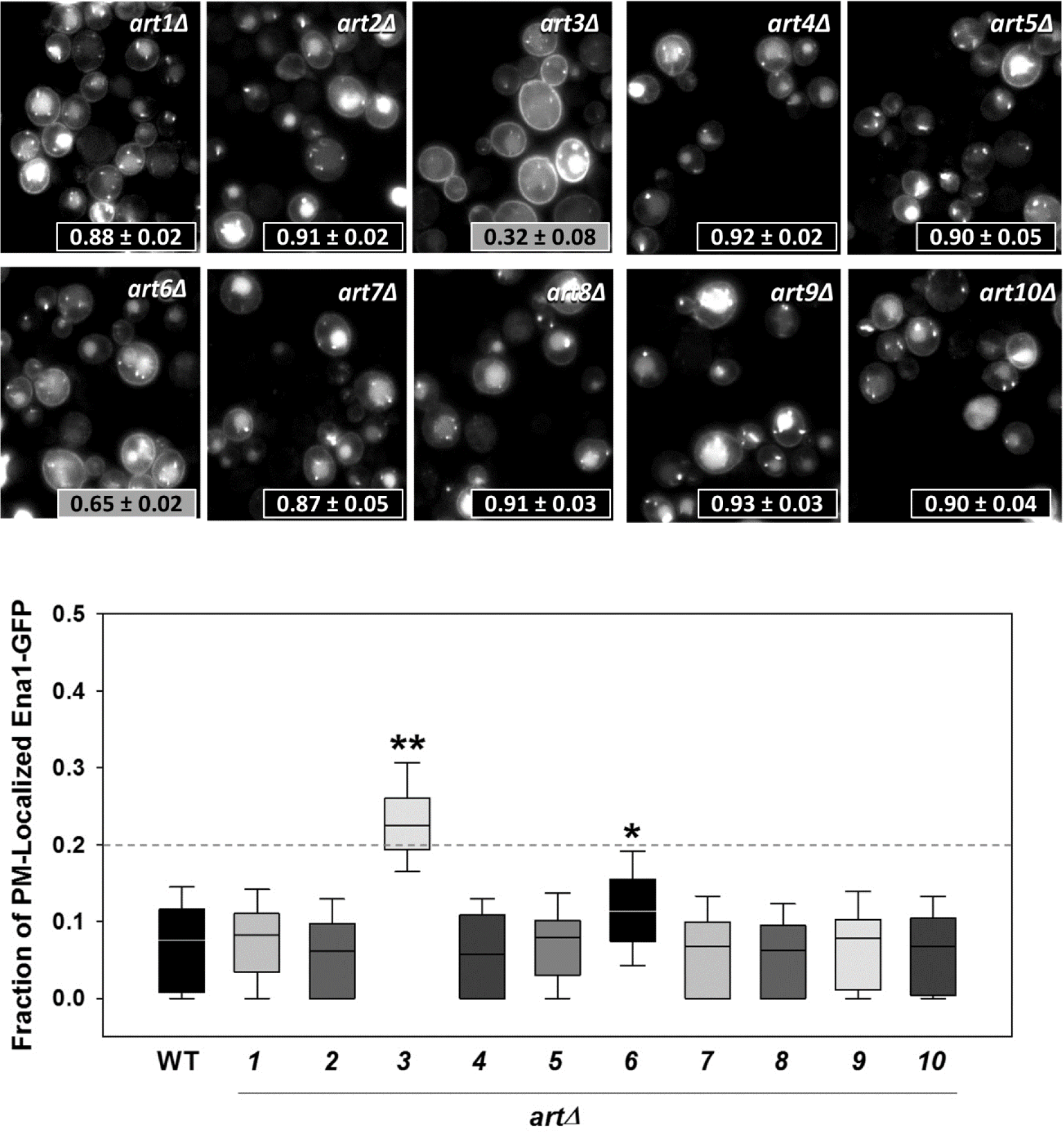
A complete collection of *ART* single deletes strains in BY4742 background expressing Ena1-GFP were imaged as described in Materials and methods. IL values and relative Ena1-GFP PM accumulation for the indicated strains was estimated and statistically analyzed with respect to the corresponding WT strain as in Fig. 1. **: p<< αC=0.001; *: p< αC=0.001—Wilcoxon’s test.

**Supplemental Fig.4.**
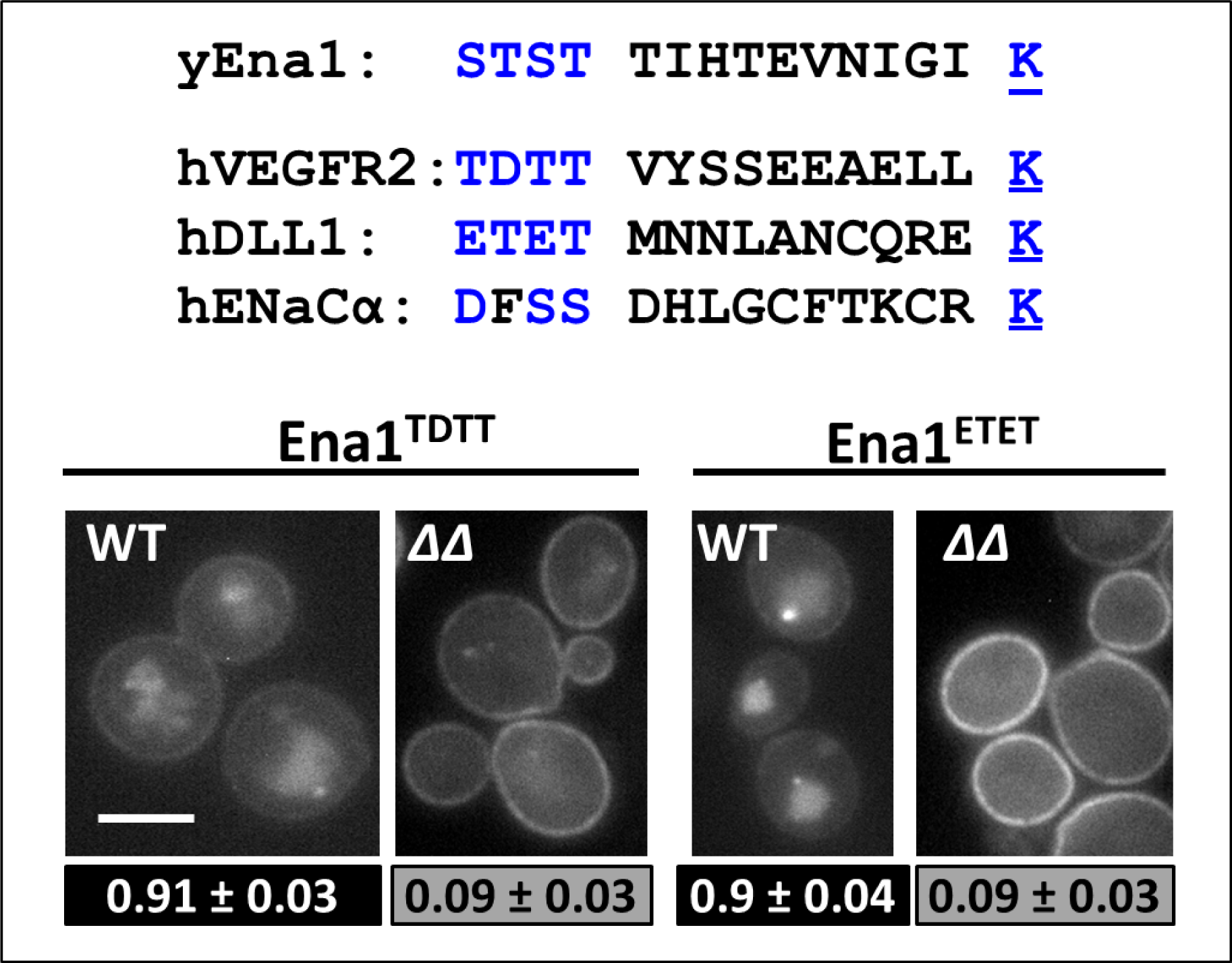
*Upper panel*: Alignment shows “STK”-like sequences (with conserved spacing from the transmembrane domain and among the ST-patch to K distance) from known mammalian epsin cargoes as compared to yeast Ena1. Chimeric proteins were created replacing Ena1 WT S^1076^TST^1079^ sequence for the hVEGFR2 TDTT (Ena1^TDTT^) and hDLL1 ETET (Ena1^ETET^) that in conjunction with K^1090^ recreated the mammalian STK motifs. y: yeast; h: human. *Lower panel*: Intracellular localization of Ena1^TDTT^-GFP (left) and Ena1^ETET^ (right) was analyzed in WT and *ΔΔ* cells. IL indexes were estimated and analyzed as described in Fig. 1. Representative images are also included. Scale bar: 5μm.

**Supplemental Fig.5.**
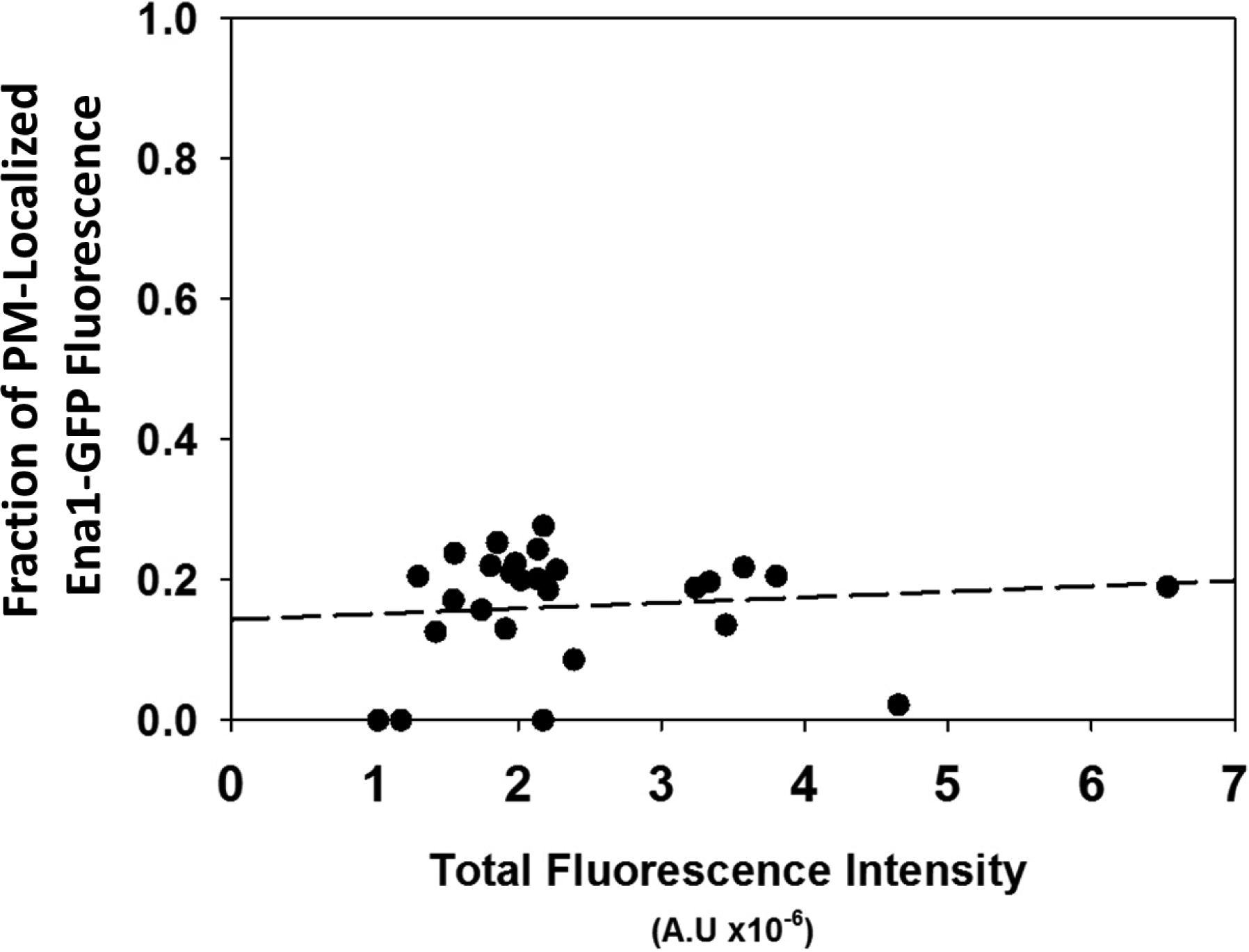
The fraction of PM-associated (P/T—see *Materials and methods*) Ena1-GFP fluorescence as a function of total cellular fluorescence intensity. Although P/T values displayed typical biological variability, their median range remained fairly constant only varying between extrapolated values of 0.14 to 0.19 (the latter representing high over-expressors—which were excluded from the current study)

**Supplemental Table I.**
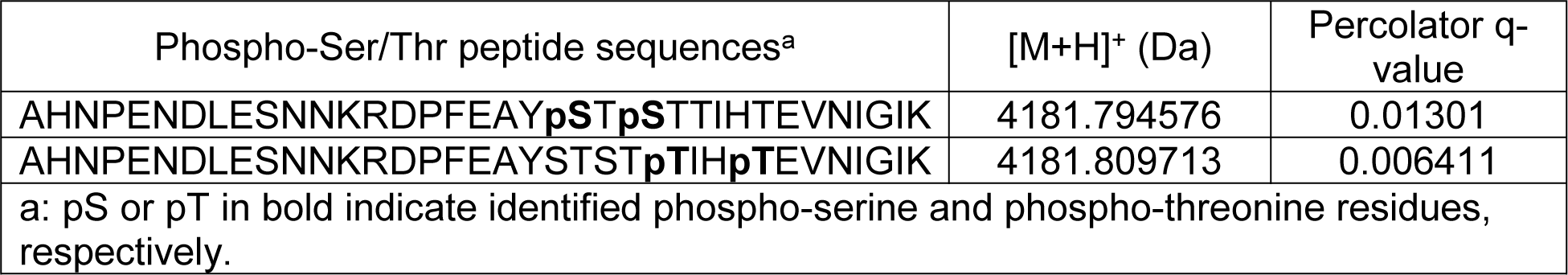
Ena1 ST-patch phosphopeptides identified by mass spectrometry

Ena1 ST-patch phosphopeptides identified by mass spectrometry
| Phospho-Ser/Thr peptide sequences<sup>a</sup> | [M+H]<sup>+</sup> (Da) | Percolator q-value |
| --- | --- | --- |
| AHNPENDLESNNKRDPFEAY**pSTp**STTIHTEVNIGIK | 4181.794576 | 0.01301 |
| AHNPENDLESNNKRDPFEAYSTST**pTIHp**TEVNIGIK | 4181.809713 | 0.006411 |
| a: pS or pT in bold indicate identified phospho-serine and phospho-threonine residues, respectively. |  |  |

**Supplemental Table II:** Plasmids used in this study

| **Name** | **Description** | **Source** |
| --- | --- | --- |
| pUG35 Ena1-GFP | *MET25::ENA1::YEGFP<sup>a</sup> CEN URA3* | (7) |
| pJLU34 (Ste3-GFP) | *STE3::GFP CEN URA3* | (8) |
| Ena1-GFP (LEU2) | *MET25::ENA1::YEGFP CEN LEU2* | This study |
| Ent1.317 | *ENT1 CEN LYS2* | Wendland lab, JHU |
| ENTH1.317 | *ENT1<sup>1-154</sup> CEN LYS2* | This study |
| ΔENTH1.317 | *ENT1<sup>145-454</sup> CEN LYS2* | This study |
| Ent1<sup>Um</sup>.317 | *ENT1<sup>S177D S201D</sup> CEN LYS2* | This study |
| Ent1<sup>1-235</sup>.317 | *ENT1<sup>1-235</sup>::HAX3 CEN LYS2* | This study |
| Ent2.317 | *ENT2 CEN LYS2* | Wendlandlab, JHU |
| ΔENTH2.317 | *ENT2<sup>148-613</sup> CEN LYS2* | This study |
| Ent2<sup>Um</sup>.317 | *ENT2<sup>S187D S218D</sup> CEN LYS2* | This study |
| Ena1<sup>1-1071</sup>-GFP | *MET25::ENA1<sup>Δ(1072-1091)</sup>::YEGFP CEN URA3* | This study |
| Ena1<sup>1-1075</sup>-GFP | *MET25::ENA1<sup>Δ(1076-1091)</sup>::YEGFP CEN URA3* | This study |
| Ena1<sup>1-1079</sup>-GFP | *MET25::ENA1<sup>Δ(1080-1091)</sup>::YEGFP CEN URA3* | This study |
| Ena1<sup>1-1083</sup>-GFP | *MET25::ENA1<sup>Δ(1084-1091)</sup>::YEGFP CEN URA3* | This study |
| Ena1<sup>1-1087</sup>-GFP | *MET25::ENA1<sup>Δ(1088-1091)</sup>::YEGFP CEN URA3* | This study |
| Ena1<sup>G1088A</sup>-GFP | *MET25::ENA1<sup>G1088A</sup>::YEGFP CEN URA3* | This study |
| Ena1<sup>I1089A</sup>-GFP | *MET25::ENA1<sup>I1089A</sup>::YEGFP CEN URA3* | This study |
| Ena1<sup>Q1091A</sup>-GFP | *MET25::ENA1<sup>Q1091A</sup>::YEGFP CEN URA3* | This study |
| Ena1<sup>K1090R</sup>-GFP | *MET25::ENA1<sup>K1090R</sup>::YEGFP CEN URA3* | This study |
| Ena1<sup>K1090R</sup>-Ub-GFP | *MET25::ENA1<sup>K1090R</sup>-Ub<sup>K48R, ΔG75, ΔG76</sup>::YEGFP CEN URA3<sup>b</sup>* | This study |
| Ena1<sup>(1088-1091)G</sup>-GFP | *MET25::ENA1<sup>I1089G K1090G Q1091G</sup>::YEGFP CEN URA3* | This study |
| Ena1<sup>(1084-1087)G</sup>-GFP | *MET25::ENA1<sup>E1084G V1085G N1086GI1087G</sup>::YEGFP CEN URA3* | This study |
| Ena1<sup>(1080-1083)G</sup>-GFP | *MET25::ENA1<sup>T1080G I1081G H1082G T1083G</sup>::YEGFP CEN URA3* | This study |
| Ena1<sup>(1076-1079)G</sup>-GFP | *MET25::ENA1<sup>S1076G T1077GS1078G T1079G</sup>::YEGFP CEN URA3* | This study |
| Ena1<sup>(1072-1075)G</sup>-GFP | *MET25::ENA1<sup>F1072G E1073GA1074G Y1075G</sup>::YEGFP CEN URA3* | This study |
| Ena1<sup>TSTSS</sup>-GFP | *MET25::ENA1<sup>S1076T T1077S S1078T T1079S T1080S</sup>::YEGFP CEN URA3* | This study |
| Ena1<sup>DGGG</sup>-GFP | *MET25::ENA1<sup>S1076D T1077GS1078G T1079G</sup>::YEGFP CEN URA3* | This study |
| Ena1<sup>GDGG</sup>-GFP | *MET25::ENA1<sup>S1076G T1077DS1078G T1079G</sup>::YEGFP CEN URA3* | This study |
| Ena1<sup>GGDG</sup>-GFP | *MET25::ENA1<sup>S1076G T1077GS1078D T1079G</sup>::YEGFP CEN URA3* | This study |
| Ena1<sup>GGGD</sup>-GFP | *MET25::ENA1<sup>S1076G T1077GS1078G T1079D</sup>::YEGFP CEN URA3* | This study |
| Ena1<sup>DDGG</sup>-GFP | *MET25::ENA1<sup>S1076D T1077DS1078G T1079G</sup>::YEGFP CEN URA3* | This study |
| Ena1<sup>DGDG</sup>-GFP | *MET25::ENA1<sup>S1076D T1077GS1078D T1079G</sup>::YEGFP CEN URA3* | This study |
| Ena1<sup>DGGD</sup>-GFP | *MET25::ENA1<sup>S1076D T1077GS1078G T1079D</sup>::YEGFP CEN URA3* | This study |
| Ena1<sup>GDDG</sup>-GFP | *MET25::ENA1<sup>S1076G T1077DS1078D T1079G</sup>::YEGFP CEN URA3* | This study |
| Ena1<sup>GDGD</sup>-GFP | *MET25::ENA1<sup>S1076G T1077DS1078G T1079D</sup>::YEGFP CEN URA3* | This study |
| Ena1<sup>GGDD</sup>-GFP | *MET25::ENA1<sup>S1076G T1077GS1078D T1079D</sup>::YEGFP CEN URA3* | This study |
| Ena1<sup>AAST</sup>-GFP | *MET25::ENA1<sup>S1076A T1077A</sup>::YEGFP CEN URA3* | This study |
| Ena1<sup>ATAT</sup>-GFP | *MET25::ENA1<sup>S1076A S1078A</sup>::YEGFP CEN URA3* | This study |
| Ena1<sup>ATSA</sup>-GFP | *MET25::ENA1<sup>S1076A T1079A</sup>::YEGFP CEN URA3* | This study |
| Ena1<sup>SAAT</sup>-GFP | *MET25::ENA1<sup>T1077A S1078A</sup>::YEGFP CEN URA3* | This study |
| Ena1<sup>SASA</sup>-GFP | *MET25::ENA1<sup>T1077A T1079A</sup>::YEGFP CEN URA3* | This study |
| Ena1<sup>STAA</sup>-GFP | *MET25::ENA1<sup>S1078A T1079A</sup>::YEGFP CEN URA3* | This study |
| Ena1<sup>ATST</sup> -GFP | *MET25::ENA1<sup>S1076A</sup> ::YEGFP CEN URA3* | This study |
| Ena1<sup>DDST</sup> -GFP | *MET25::ENA1<sup>S1076D T1077D</sup> ::YEGFP CEN URA3* | This study |
| Ena1<sup>AAST</sup> -Ub -GFP | *MET25::ENA1<sup>S1076A S1077A</sup> - Ub<sup>K48R, ΔG75, ΔG76</sup> ::YEGFP CEN URA3<sup>a</sup>* | This study |
| Ena1<sup>DDST, K1090R</sup> -GFP | *MET25::ENA1<sup>S1076D T1077D K1090R</sup> ::YEGFP CEN URA3* | This study |
| Ena1<sup>DDST, K1090R</sup> -Ub-GFP | *MET25::ENA1<sup>S1076D T1077D K1090R</sup> -Ub<sup>K48R, Δ(G75-G76)</sup> ::YEGFP CEN URA3<sup>a</sup>* | This study |
| Ena1<sup>TDTT</sup> -GFP | *MET25::ENA1<sup>S1076T T1077D S1078T</sup> ::YEGFP CEN URA3* | This study |
| Ena1<sup>ETET</sup> -GFP | *MET25::ENA1<sup>S1076E S1078E</sup> ::YEGFP CEN URA3* | This study |
| Ena1<sup>dupl(1066-1070)</sup> -GFP | *MET25::ENA1::YEGFP with N<sup>1066</sup>NRRD<sup>1070</sup> duplicated CEN URA3* | This study |
| Ena1<sup>dupl(1081-1085)</sup> -GFP | *MET25::ENA1::YEGFP with I<sup>1081</sup>HTEV<sup>1085</sup> duplicated CEN URA3* | This study |
| Ena1<sup>Δ(1083-1087)</sup> -GFP | *MET25::ENA1<sup>Δ(1083-1087)</sup> ::YEGFP CEN URA3* | This study |
| Ena1<sup>Δ(1058-1091)</sup> -GFP | *MET25::ENA1<sup>Δ(1058-1091)</sup> ::YEGFP CEN URA3* | This study |
| GFP-Tre1 | *TPI1::GFP::TRE1 in Ycplac33, CEN URA3* | (9) |
| pLJ721(Yck1) | *YCK1 2μ URA3* | (10) |
| pLS2.3 (Yck2) | *YCK2 2μ URA3* | (10) |
| Yck1-His<sub>6</sub> | *GAL10::YCK1<sup>1-536</sup> -His<sub>6</sub> 2μ URA3* | This study |
| Yck2-His<sub>6</sub> | *GAL1::YCK1<sup>1-546</sup> -His<sub>6</sub> 2μ URA3* | This study |
| Art3. 425 | *ART3 2μ, LEU2* | (11) |
| HA-ΔENTH1. 426 | *MET25::HAX3::ENT1<sup>149-454</sup> 2μ, URA3* | This study |
| HA-ΔENTH1. 317 | *ENT1<sup>149-454</sup> ::HAX3 CEN LYS2* | This study |
| His<sub>6</sub>-Ent1 | pET28a His<sub>6</sub>-Ent1 | (5) |
| His<sub>6</sub>-Ent2 | pET28c His<sub>6</sub>-Ent2 | Lab collection |
| Ub-GST | pET3a Ub-GST | kindly provided by the late Cecile Pickart (Johns Hopkins) |
| Ena1<sup>K1090R</sup> -Ub-GST | pET3a Ena1<sup>1076-1091, K1090R</sup> - Ub-GST | This study |
| DDST-Ub-GST | pET3a Ena1<sup>1076-1091, S1076DT, 1077D</sup> - Ub-GST | This study |
| GST | pGEX-4T-1 | Lab collection |
| pRS 425 | *2μ, LEU2* | (12) |
| pRS 317 | *CEN, LYS* | Lab collection |
| pRS 426 | *2μ, URA3* | Lab collection |

a: Enhanced Green Fluorescence Protein with yeast codon optimization.
b: The ubiquitin moiety fused to the constructs was Ub<sup>K48R, ΔG75, ΔG76</sup>, where K48R prevents formation of K48 linked polyubiquitin chains that have been reported to primarily signal degradation (13), and elimination of two C- terminal glycine residues prevents recognition and cleavage by deubiquitinating enzymes (14).

**Supplemental Table III:**
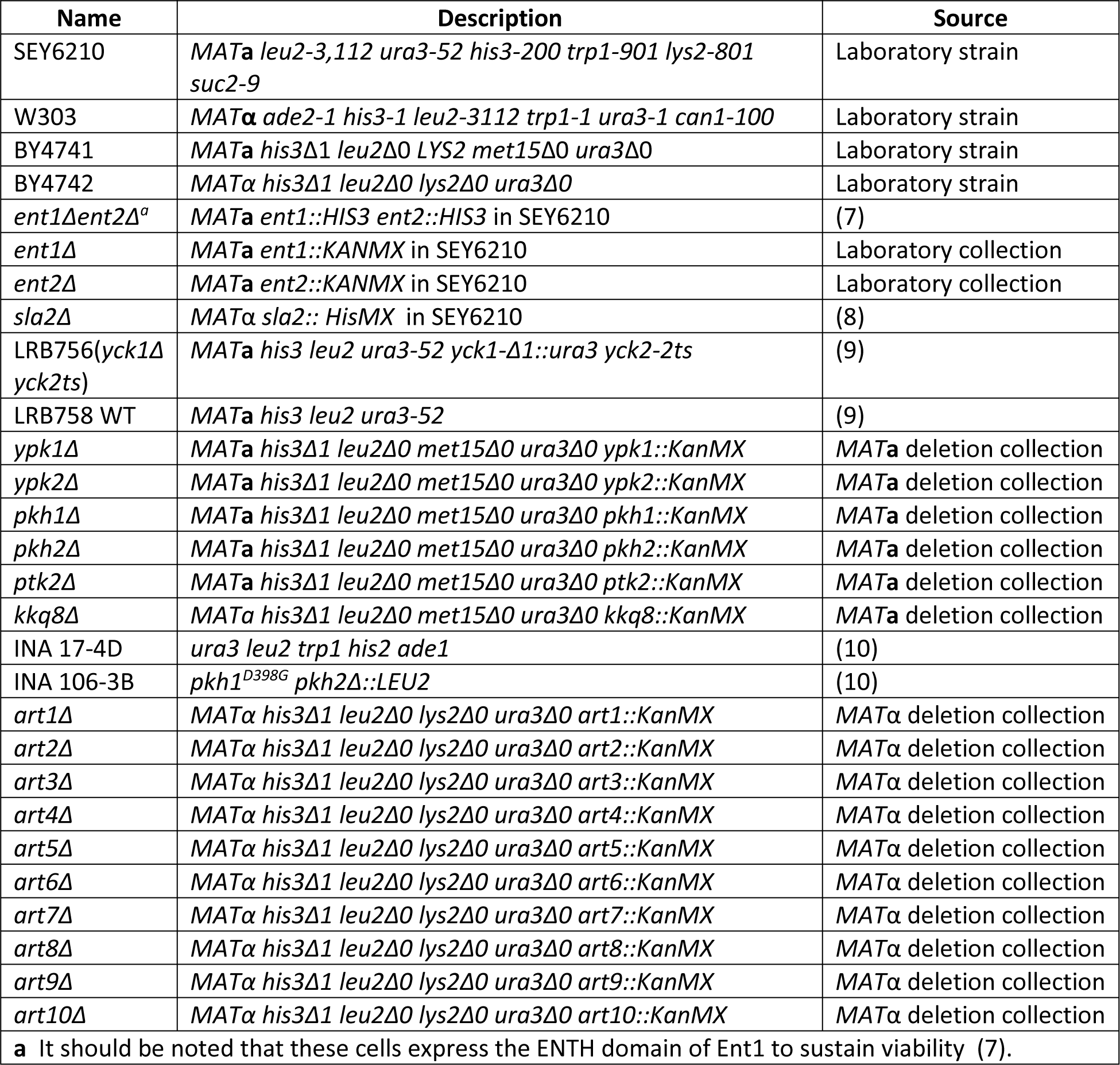
Strains used in this study

